# Differential aberrant structural synaptic plasticity in axons and dendrites ahead of their degeneration in tauopathy

**DOI:** 10.1101/2020.04.29.067629

**Authors:** Johanna S. Jackson, James D. Johnson, Soraya Meftah, Tracey K Murray, Zeshan Ahmed, Matteo Fasiolo, Michael L. Hutton, John T.R. Isaac, Michael J. O’Neill, Michael C. Ashby

**Affiliations:** Eli Lilly & Co. Ltd., Erl Wood Manor, Windlesham, Surrey GU20 6PH, UK; School of Physiology, Pharmacology and Neuroscience, University of Bristol, Biomedical Sciences Building, University Walk, Bristol BS8 1TD, UK; UK Dementia Research Institute at Imperial College, Department of Brain Sciences, Imperial College London, London, UK; John Edward Porter Neuroscience Research Center (Building 35), National Institutes of Health, 35 Convent Drive, Bethesda, MD 20814, USA; School of Mathematics, University of Bristol, Fry Building, Woodland Road, Bristol BS8 1UG, UK; Neuroscience at the J&J London Innovation Centre, One Chapel Place, London, W1G 0BG, UK; AbbVie Deutschland GmbH & Co. K.G., Ludwigshafen, Germany

## Abstract

Neurodegeneration driven by aberrant tau is a key feature of many dementias. Pathological stages of tauopathy are characterised by reduced synapse density and altered synapse function. Furthermore, changes in synaptic plasticity have been documented in the early stages of tauopathy suggesting that they may be a driver of later pathology. However, it remains unclear if synapse plasticity is specifically linked to the degeneration of neurons. This is partly because, in progressive dementias, pathology can vary widely from cell-to-cell along the prolonged disease time-course. To overcome this variability, we have taken a longitudinal experimental approach to track individual neurons through the progression of neurodegenerative tauopathy. Using repeated *in vivo* 2-photon imaging in rTg4510 transgenic mice, we have measured structural plasticity of presynaptic terminaux boutons and postsynaptic spines on individual axons and dendrites over long periods of time. By following individual neurons, we have measured synapse density across the neuronal population and tracked changes in synapse turnover in each neuron. We found that tauopathy drives a reduction in density of both presynaptic and postsynaptic structures and that this is partially driven by degeneration of individual axons and dendrites that are spread widely across the disease time-course. Both synaptic loss and neuronal degeneration was ameliorated by reduction in expression of the aberrant P301L transgene, but only if that reduction was initiated early in disease progression. Notably, neurite degeneration was preceded by alterations in synapse turnover that contrasted in axons and dendrites. In dendrites destined to die, there was a dramatic loss of spines in the week immediately before degeneration. In contrast, axonal degeneration was preceded by a progressive attenuation of presynaptic turnover that started many weeks before axon disappearance. Therefore, changes in synapse plasticity are harbingers of degeneration of individual neurites that occur at differing stages of tau-driven neurodegenerative disease, suggesting a cell or neurite autonomous process. Furthermore, the links between synapse plasticity and degeneration are distinct in axonal and dendritic compartments.

**Key findings:**

- Tauopathy driven by tau P301L in rTg4510 mice causes a progressive decrease in density of presynaptic terminaux boutons and postsynaptic dendritic spines in cortical excitatory neurons.
- Longitudinal imaging of individual axons and dendrites shows that there is a huge diversity of effects at varying times in different cells.
- Decreases in overall synapse density are driven partly, but not exclusively, by degeneration of dendrites and axons that are distributed widely across the time-course of disease.
- Suppression of pathological P301L tau expression can ameliorate accumulation of tau pathology, synapse loss and neurodegeneration, but only if administered early in disease progression.
- Neurite degeneration is preceded by aberrant structural synaptic plasticity in a cell-specific way that is markedly different in dendrites and axons.
- Degeneration of dendrites is immediately preceded by dramatic loss of dendritic spines.
- Axonal loss is characterised by a progressive attenuation of presynaptic bouton plasticity that starts months before degeneration.

## Introduction

Decreased synapse density is a cardinal feature of neurodegenerative dementia (Scheff et al., 2006; Spires-Jones and Hyman, 2014). Furthermore, dysfunctional neurotransmission and changes in synaptic plasticity have been documented in many animal models of dementia, leading to the idea of synaptic normalisation as a potential therapeutic target (Forner et al., 2017; Herms and Dorostkar, 2016; Jackson et al., 2019). However, the links between changes in synaptic plasticity, synapse loss and neurodegeneration are still poorly understood.

In tauopathy-related dementias, such as Alzheimer’s Disease (AD), increasingly aberrant and hyperphosphorylated tau is associated with synapse loss, neuronal death and the cognitive symptoms that are observed (Gendron and Petrucelli, 2009; Nelson et al., 2012). Aberrant tau in various forms is almost ubiquitously reported to cause overall reduction in synapse number, including observations in the rTg4510 mouse model, which expresses a hyperphosphorylated P301L version of tau (Jackson et al., 2017; Kopeikina et al., 2013). Both pre- and postsynaptic impairments driven by aberrant tau have been documented using multiple methods across a range of disease timepoints. At presynaptic sites, neurotransmitter release probability is reduced in human tau-expressing mice (Polydoro et al., 2009). This impairment is potentially mediated by the N-terminal of tau directly associating with pre-synaptic vesicles, which causes impairment in prolonged synaptic release and synaptic vesicle motility in fly and rat neurons (Zhou et al., 2017). Additionally, P301L tau induces changes in presynaptic vesicular glutamate transporters and glutamate transporter 1 (GLT-1), resulting in increased glutamate release and decreased extracellular clearance (Hunsberger et al., 2015). Presynaptic electrophysiological deficits have also been observed in transgenic mice which express the P301L tau mutation (Polydoro et al., 2009, 2014). In contrast, overexpressed human TauP301L has been shown to accumulate at presynaptic sites without causing any cognitive disruption (Harris et al., 2012). Although tau is predominantly localised to axons in healthy neurons, elevated levels of hyperphosphorylated tau is found in the somatodendritic compartment, including perisynaptic areas (Hoover et al., 2010; Ittner et al., 2010). Indeed, it has been suggested that hyperphosphorylated tau may actually be concentrated at postsynaptic locations (Tai et al., 2012), perhaps due to its trans-synaptic spread that can occur early in the disease (Pickett et al., 2017). As such, many studies investigating synapse loss and dysfunction have focused on the postsynaptic changes associated with aberrant tau (Ittner and Ittner, 2018). Postsynaptic dysfunction can be driven by missorting of tau into the postsynaptic region along with the Src kinase, Fyn (Ittner et al., 2010). Fyn alters NMDAR phosphorylation and its interaction with the postsynaptic density protein 95 (PSD-95). This over-stabilises NMDARs at the postsynaptic site, resulting in excessive calcium influx and damage through excitotoxicity (Ittner et al., 2010; Mondragón-Rodríguez et al., 2012). There is also substantial evidence that aberrant tau can affect synaptic plasticity. Synaptic long-term potentiation (LTP) is reduced in transgenic mouse lines expressing mutated forms of tau (Rosenmann et al., 2008; Yoshiyama et al., 2007). Reduced size and prevalence of mature “mushroom” dendritic spines suggests a weakening of synapses driven in mouse models of tauopathy (Crimins et al., 2012; Jackson et al., 2017). Furthermore, the structural plasticity underlying addition and removal of synapses is impacted in the rTg4510 mouse (Jackson et al., 2017). Given the central role of NMDARs in induction of several forms of synaptic plasticity, it is tempting to link tau-induced changes in synapse plasticity to spine loss and subsequent neurodegeneration, but it is unclear how, when and where in the cell these processes might coincide (Ittner and Ittner, 2018; Miyamoto et al., 2017; Tackenberg and Brandt, 2009).

Several studies have identified synaptic changes that occur ahead of the onset of classic histopathological markers and frank neurodegeneration, suggesting a causal role in progression of tauopathy (Menkes-Caspi et al., 2015; Rocher et al., 2010; Yoshiyama et al., 2007; Zhou et al., 2017). However, there are even more reports of synaptic dysfunction and/or aberrant plasticity during later, neurodegenerative phases of tauopathy (Booth et al., 2016; Crimins et al., 2012). Some of these late changes do not always mirror those found early in disease, even in similar brain areas. It is perhaps unsurprising that synaptic alterations may go through different phases as tauopathy progresses, but these contradictions highlight the lack of understanding of the specificity and timing of pre- and post-synaptic changes in relation to the pathology consuming the parent neurons.

In this study, we have used longitudinal *in vivo* two-photon imaging over extended periods to track synaptic alterations and degeneration of neuronal processes across the time-course of tauopathy in rTg4510 mice. Tracking individual axons and dendrites at frequent intervals across long periods of time allowed us to capture the diversity of neuronal changes related to different stages of progressive tauopathy, including the moment of neurite degeneration. We have identified alterations in synapse turnover that manifest in the weeks before neurite degeneration. These changes are strikingly different in pre- and post-synaptic compartments in both their effects and in their time-course.

## Results

### The time-course of synapse loss in rTg4510 mice

To characterise the dynamics of synapse turnover and degeneration across the time-course of tauopathy, we used *in vivo* 2-photon microscopy to repeatedly image axonal and dendritic structure of cortical pyramidal neurons in rTg4510 mice. We imaged from early, pre-pathological stages through to ages associated with overt signs of neurodegenerative disease. To enable chronic measurements, we implanted cranial windows in rTg4510 mice and wild-type (WT) littermates having transduced the underlying somatosensory cortex with AAV that drives expression of GFP in excitatory neurons. Following a post-surgical recovery period, GFP-expressing dendritic and axonal branches that ramified in layer 1 were imaged in head-fixed anaesthetised mice. To ensure coverage of a large portion of the progressive development of pathology, we imaged animals in four batches that were staggered in their age at the start of imaging (20, 24, 28 and 32 weeks old). Within these batches, animals were imaged weekly for up to 26 weeks (median number of imaging sessions was 16). This experimental design produced longitudinal data that spanned from approximately 4.5 months old, when there is little cortical atrophy in rTg4510 mice, to 12 months old, when these mice have substantial cortical loss and display dramatic neurological deficits (Ramsden et al., 2005; Holmes et al., 2016).

First, we measured alterations in synapse density in labelled cells by identifying and counting dendritic spines and axonal terminaux boutons (TBs) in each imaging session (Figures 1&2). Dendrites were distinguished by a relatively straight and wide shaft with spines protruding that had characteristic bulbous heads and short necks (Holtmaat et al., 2009)(Figure 1A). In contrast, axons had a thinner shaft with a greater tortuosity (Holtmaat et al., 2009)(Figure 1A & 2A). Most axons were studded with protruding terminaux boutons (TBs) that varied dramatically in shape but tended to have long necks, as well as occasional *en passant* swellings (Figure 1A & 2A). In total, we imaged 114 dendrites across 1688 image stacks (Figure 1) and 106 axons across 1382 image stacks (Figure 2) from 20 WT and 20 rTg4510 animals.

**Figure 1.**
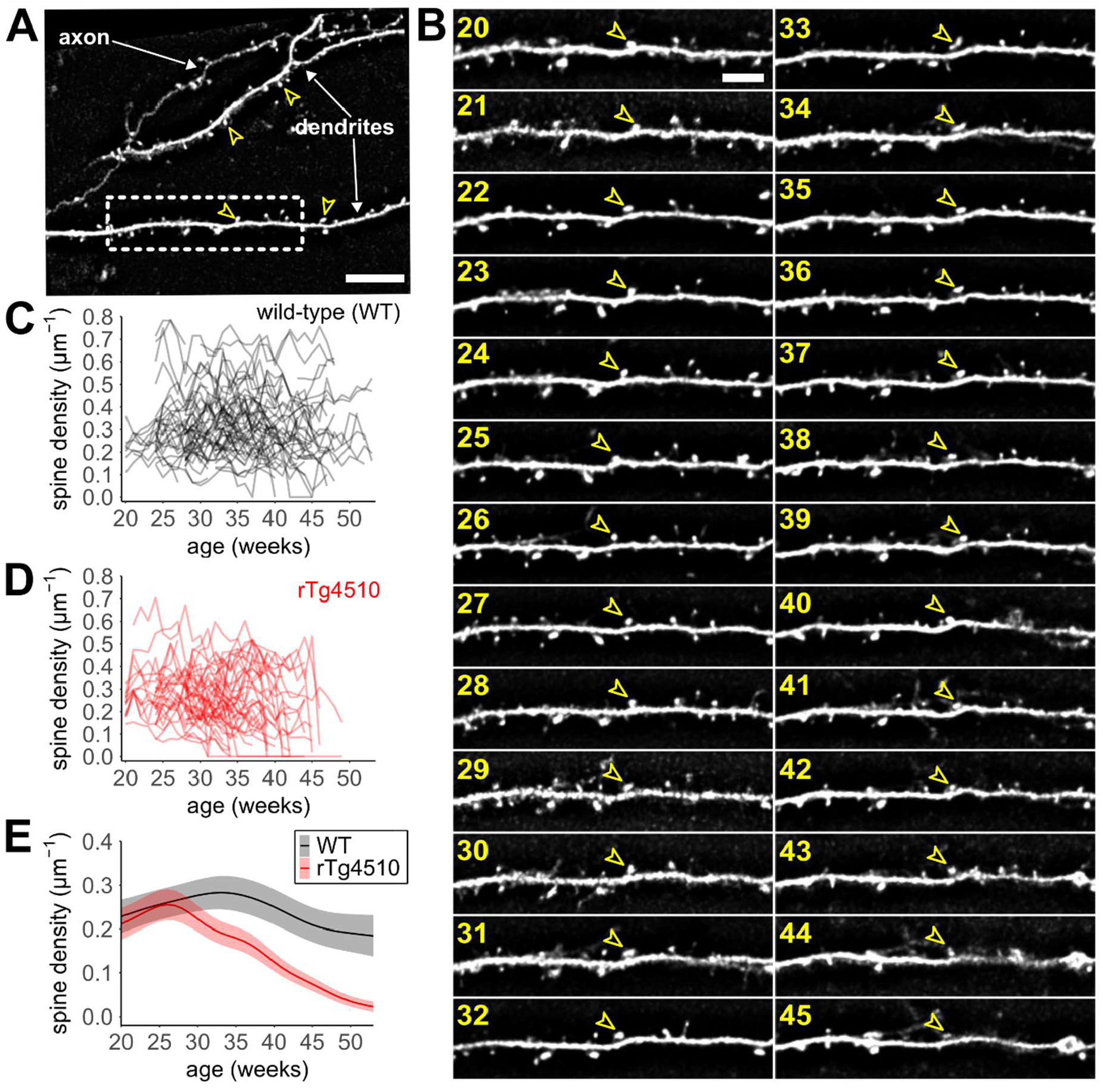
Longitudinal tracking of dendritic spine loss in rTg4510 mice. **(A)** Typical 2-photon field of view of somatosensory cortex containing sparsely-labelled dendrites. Example dendritic spines marked with empty arrowheads. Note that there is also an axon in the upper left of the filed of view. Scale bar, 2 μm. **(B)** Example of repeated weekly imaging of the same dendritic branch (from dashed box in (A)) across periods of time (age in weeks shown in yellow). Individual spines were identified in each imaging session (example shown by arrowhead). Scale bar, 5 μm. **(C)** Density of spines over time for each of 60 dendrites tracked in 20 wild-type (WT) mice. **(D)** Density of spines over time for each of 54 dendrites tracked in 20 rTg4510 tauopathy mice. **(E)** Predictions from a GAMM based on data in (B) and (C) modelling changes in overall spine density across the population over time (shaded area represents 95% confidence limits of model estimate shown by line).

**Figure 2.**
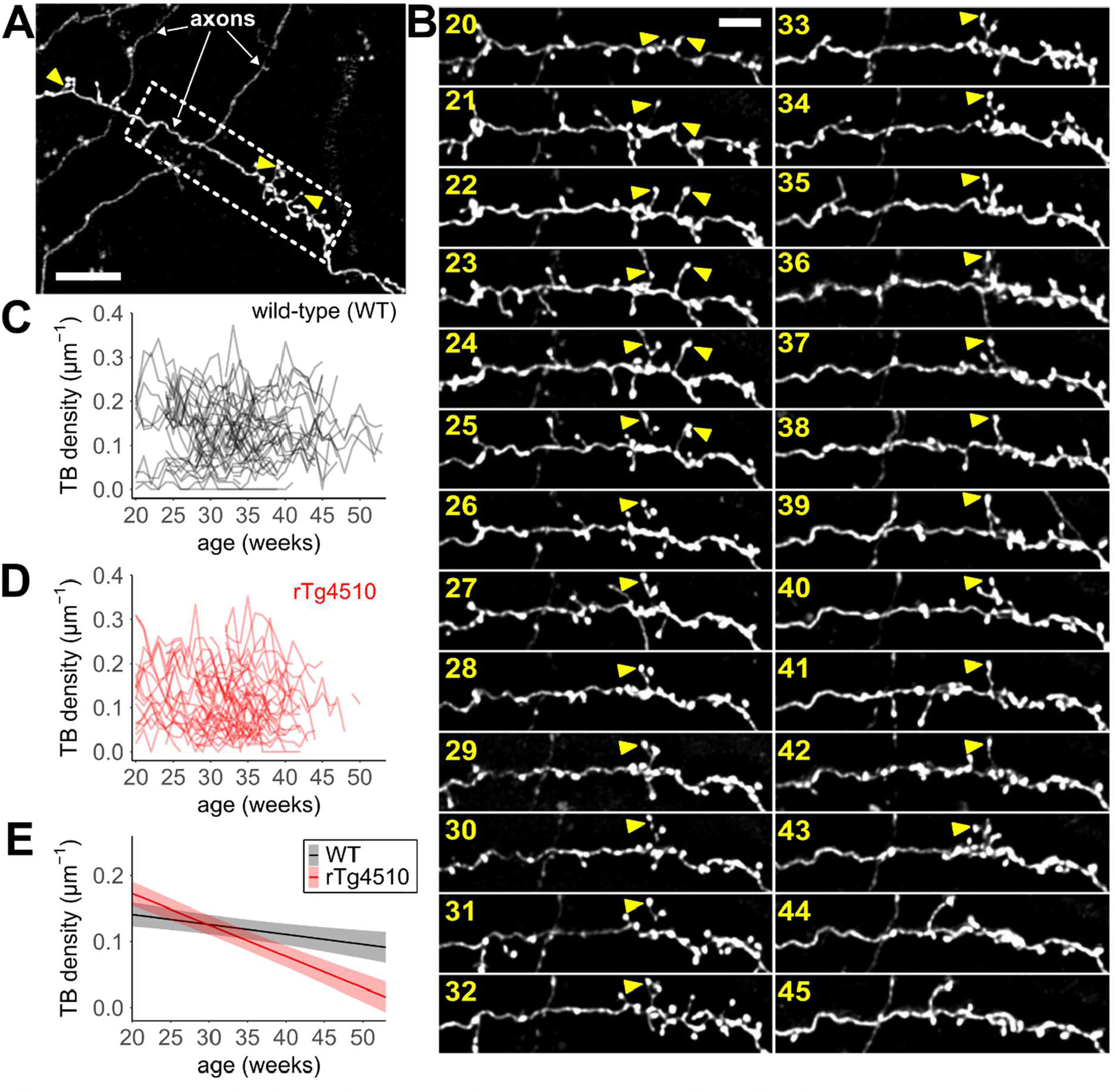
Longitudinal tracking of terminaux boutons in rTg4510 mice. **(A)** Typical 2-photon field of view of somatosensory cortex containing sparsely-labelled axons. Example terminaux boutons (TBs) are marked with arrowheads. Scale bar, 20 μm. **(B)** Example of repeated weekly imaging of the same axonal branch (from dashed box in (A)) across periods of time (age in weeks shown in yellow). Individual TBs were identified in each imaging session (examples shown by arrowhead). Scale bar, 5 μm. **(C)** Density of TBs over time for each of 51 axons tracked in 20 wild-type (WT) mice. **(D)** Density of spines over time for each of 55 axons tracked in 20 rTg4510 tauopathy mice. **(E)** Predictions from a GAMM based on data in (C) and (D) modelling changes in overall TB density across the population over time (shaded area represents 95% confidence limits of model estimate shown by line).

Within each imaged dendrite, we could readily identify many spines that persisted across long periods of time as well as others that appeared or disappeared from one week to the next (Figure 1B). The longitudinal imaging of these structural dynamics allowed us to determine the density of spines on each dendrite across time (Figure 1C&D; n=60 WT, 54 rTg4510 dendrites from 20 mice). We observed quite divergent dynamics in spine density in individual dendrites (Figure 1C&D). This was particularly the case in rTg4510 animals in which some neurites showed dramatic losses of spines or TBs, whereas others had relatively stable density even at older ages (Figure 1D). To estimate the overall effect of genotype on the dynamics of spine density across the entire imaged population we fitted Generalised Additive Mixed Models (GAMMs) to these data. GAMMs have the major advantage of not enforcing predetermined assumptions about the trajectory of the data while also accounting for longitudinality (including missing timepoints) and hierarchical experimental design (i.e. multiple neurons within each animal, see Methods for further description)(van Rij et al., 2019; Shadish et al., 2014; Wood, 2017). Overall, the GAMM fit the dataset well, explaining 83.9% of deviance for dendritic spine density data (Rsq = 0.85). In line with previous data, population spine density was relatively stable over time in WT animals (Figure 1E; Grutzendler et al., 2002; Jackson et al., 2017). In contrast, the overall density of postsynaptic spines progressively decreased with age in rTg4510s (Figure 1E). In the early stages of our imaging, which corresponds to largely pre-degenerative phases of the disease, WT and rTg4510 dendritic spine density was similarly stable around ∼0.2μm^-1^ (Figure 1E). Between 25-30 weeks of age, the trajectory of WT and rTg4510 diverge, with WT remaining relatively stable for the next few months of imaging, whereas there is a continual decline in the spine density of rTg4510s across this period. By the end of our imaging period, at ∼50 weeks old, the majority of, but not all, dendrites in rTg4510 animals either have very few spines or have disappeared altogether. Genotype has a significant effect on model estimates (Wald test, p = 0.025 for genotype as fixed factor) as suggested by the diverging trajectories for each WT and rTg4510 dendrites. We further tested the influence of each factor in the GAMM by comparing to an analogous model lacking the factor in question. This approach showed that genotype has a major influence on the predictions as its inclusion significantly improved fit to the data (lower Akaike Information Criteria (AIC), Chi-square test, p<0.001). In contrast, variance associated with individual animals had only small effect on model predictions (Supplementary Figure 1A) and the batch had almost no effect (Supplementary Figure 1B).

In axons, we were again able to track individual TBs over long periods of time, but these protrusions were generally more dynamic than dendritic spines in their addition/removal and in their shape (Figure 2B). As with dendrites, longitudinal imaging made it apparent that changes in TB density varied dramatically in different axons (Figure 2C&D, n=51 WT, 55 rTg4510 axons from 20 mice). To assess the overall impact of these variable dynamics, we again fitted a GAMM to the WT and rTg4510 data (Figure 2E; 85.5% deviance explained, Rsq = 0.83). This analysis showed that the overall population density of TBs was ∼0.15μm^-1^ in both WT and rTg4510 animals at the youngest ages we imaged but there were differing trajectories over time for each genotype (Figure 2E). Similarly to dendritic spines, the TB density in rTg4510 animals declined over time compared to WT (Figure 2E). Effects associated with individual animals (Supplementary Figure 1E) or batch (Supplementary Figure 1F) had very little effect on model outcomes. The divergence between genotypes appeared to manifest slightly later than for dendritic spines (between 30-35 weeks, compare to Figure 1E). However, the decline in presynaptic density was similarly progressive as TB numbers fell to low values by ∼50 weeks old (Figure 2E).

### Progressive degeneration is rescued by early reduction in pathological tau expression

Tauopathy has been characterised as a runaway process in which aberrant tau can propagate pathology by promoting malfunction of previously healthy protein and cells (Mudher et al., 2017). In this type of “spreading” mechanism, progression of neuropathology becomes increasingly independent of the initial insult. This independence can be tested in rTg4510 mice, because expression of the P301L mutant tau gene is under the control of a tet-off promoter, which allows its suppression by administration of doxycycline (DOX)(Ramsden et al., 2005). Indeed, when expression is suppressed late in the rTg4510 disease time-course, reducing mutant tau levels is less effective in reducing gross pathology (Holmes et al., 2016). This does suggest a disconnect between mutant tau expression and gross neuropathology once the disease process takes root, but we do not know how and when synaptic changes are shaped by this. To understand the link between aberrant tau expression and synaptic degeneration, we continually dosed rTg4510 animals with DOX starting at varying timepoints aligned to our other imaging batches (18, 22, 26 and 30 weeks old, started at cranial window implantation).

Histopathological and qPCR analysis was performed on a subset of brains following fixation at the end of each imaging time-course to assess the impact of DOX administration on tau expression and progression of gross pathology. We confirmed that DOX administration did indeed reduce tau P301L mRNA levels, although not to WT levels (Figure 3A). Tau P301L expression in vehicle-treated rTg4510 animals was associated with a large reduction in size of the neocortex (47% reduction in area; Figure 3B), paralleling the forebrain atrophy previously described in this model (Ramsden et al., 2005). The cortical area of the DOX-treated rTg4510 animals was overall partially restored towards WT levels (38% recovery from mean rTg4510-associated decrease). However, the cortical area varied widely between different animals, from values close to WT through to dramatic atrophy that was similar to that in untreated rTg4510s (Figure 3B). We reasoned that the differing age at the start of dosing may underlie some of the variability between DOX-treated animals. Therefore, we compared the effects of DOX based on when administration was begun. The extent of tau suppression appeared to be independent of either the age at which DOX treatment was started (Figure 3C, 2-way ANOVA, F(3,38)=2.32, p=0.091), the age at the point of measurement or the total duration for which DOX was administered (Supplementary Figure 2A&B). These expression profiles suggest that DOX lowers tau P301L expression consistently across the time-course of the disease. To assess how the reduction of tau expression affected development of pathology, we measured cortical levels of PG5, which is a histological marker of hyperphosphorylated tau (Figure 3D). PG5 staining was similarly prominent across the forebrain in all untreated rTg4510s (age range at staining was 30-54 weeks; Supplementary Figure 2C) and was negligible in WT mice (Figure 3D&E). We compared the effects of DOX treatment started at the differing ages on PG5 staining. There was a significant interaction between effects of genotype and onset of DOX treatment on PG5 staining (2-way ANOVA, F(6,34)=7.25, p<0.001). Post-hoc analysis of simple main effects confirmed DOX treatment beginning at 18 weeks old significantly reduced PG5 staining compared to rTg4510s (p<0.001) to just above WT levels (Figure 3D, p=0.029). However, DOX administration at later ages (22, 26 and 30 weeks old) was less effective in reducing PG5 staining as PG5 levels in animals treated from 18 weeks were significantly lower than when treatment was started later (Figure 3D; p<0.001 compared to each later group). PG5 levels were not related to the total duration of DOX treatment or the age at perfusion (Supplementary Figure 2D). Therefore, even though mutant tau expression is reduced similarly when suppression is started at differing ages, the accumulation of hyperphosphorylated tau is poorly suppressed if DOX is not administered early. Importantly for synaptic imaging, the cranial window over the right hemisphere did not appear to affect progression of tauopathy or neuroinflammation as PG5 and IBA-1 levels were similar in both hemispheres across all genotypes (Supplementary Figure 2E&F).

**Figure 3.**
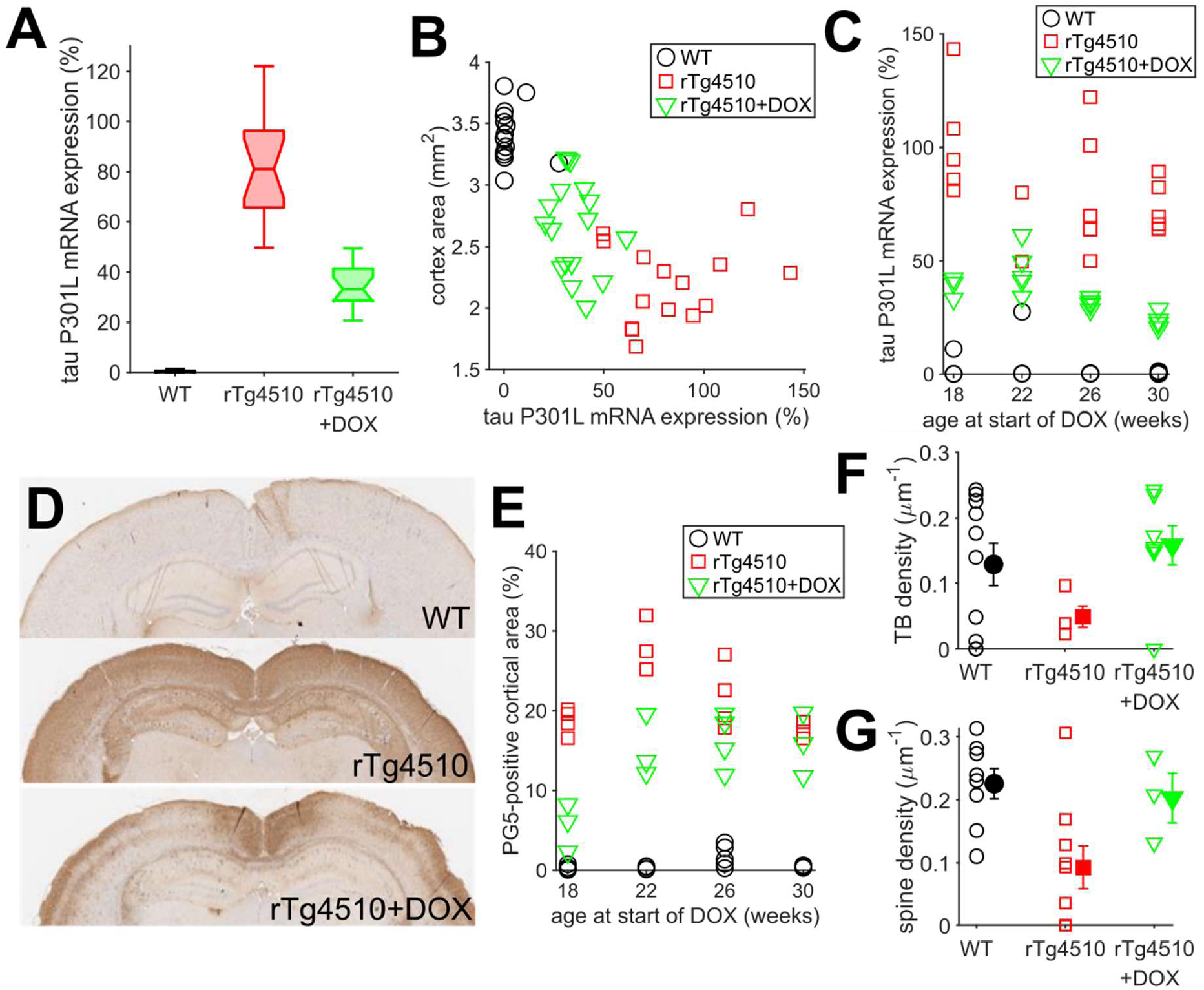
DOX treatment alleviates neuropathology if administered early. **(A)** Relative expression levels of tau P301L mRNA in brains of WT, rTg4510 and rTg4510 mice treated with DOX measured by qPCR. Central line of boxplots shows median, notches show 95% confidence intervals, whiskers show extremes of data, n = 16 WT, 17 rTg4510, 17 rTg4510+DOX animals. **(B)** Area of the neocortex is smaller in rTg4510 and those treated with DOX. **(C)** Relative tau P301L expression in rTg4510 animals treated with DOX starting at different ages, alongside age-matched WT and untreated rTg4510s. **(D)** Histological staining for PG5 in coronal brain slices containing somatosensory cortex. **(E)** Amount of cortical PG5 staining in rTg4510 animals treated with DOX starting at different ages, alongside age-matched WT and untreated rTg4510s. **(F)** Median presynaptic TB density in individual axons (empty symbols) between 35-45 weeks of ago in WT, rTg4510 and rTg4510 animals treated with DOX from 18 weeks old. DOX treatment from 18 weeks old rescues the decrease in average TB density in rTg4510 animals (filled symbols, mean+/-SEM). **(G)** Median postsynaptic spine density in individual dendrites (empty symbols) between 35-45 weeks of age in WT, rTg4510 and rTg4510 animals treated with DOX from 18 weeks old. DOX treatment from 18 weeks old rescues the decrease in average spine density in rTg4510 animals (filled symbols, mean+/-SEM).

Given its ameliorative effect on pathological markers, we wanted to assess whether DOX administration from 18 weeks also affects synapses. We therefore compared synapse density in age-matched imaging sessions from 35-45 weeks old, which is an age range where decreased synapse density is predicted (Figures 1&2). Specifically, we measured the median TB and spine density across this time period in individual axons and dendrites in WT, rTg4510 and DOX-treated (from 18 weeks old) rTg4510 animals (Figure 3F&G). We found that, overall, DOX rescued the decrease in TB density in rTg4510 animals (Figure 3F; mixed model ANOVA – animal as random factor, F(1,6.115)=8.11, p=0.029; post-hoc Tukey: WT vs rTg4510, p=0.006, rTg4510 vs DOX, p=0.001). Likewise, spine density was also decreased in rTg4510 animals compared to WT across this age range, and this decrease was rescued by early DOX treatment (Figure 3G; mixed model ANOVA– animal as random factor, F(2,17)=5.58, p=0.014; post-hoc Tukey: WT vs rTg4510, p=0.003, rTg4510 vs DOX, p=0.056). Comparison of individual axons and dendrites showed that there was considerable diversity in synapse density in all groups between different cells. This variability in synapse density in individual axons or dendrites did not appear to correlate directly with the level of tau P301L expression assessed in post-mortem brain tissue (Supplementary Figure 3). This suggests that synapse loss in rTg4510s is a rather cell-specific phenomenon. Even at seemingly similar global tau P301L levels, different neurons can range from WT synapse density to very low values or even zero in neurites that disappeared, taking their synapses with them (Figures 1BD & Supplementary Figure 3).

### Synapse loss partially relates to dendritic and axonal degeneration

The repeated weekly imaging within our 2-photon imaging dataset allowed us to document the disappearance of some axons and dendrites during the imaging timeline. In many cases there was specific loss of an individual neurite while others within the same field of view remained visible and apparently healthy (example in Figure 4C). As such, these neurite losses appeared to be specific neurodegeneration events. To assess these disappearances of individual neurites, we generated survival curves for all dendrites and axons for ∼6 months across the experiment (Figure 4A&B). During this time, there was neurite-specific loss of a substantial minority of imaged dendrites and axons in rTg4510 mice. Overall, ∼35% of dendrites (Figure 4A; n = 54 dendrites from 20 animals) and ∼14% of axons (Figure 4B; n = 55 axons from 20 animals) were lost between 2.5 and 8.5 months old in rTg4510 animals, whereas there were no neurites losses in WT mice (n = 60 dendrites and 51 axons from 20 mice). These neurite losses align with the well-documented gross neurodegenerative phenotype caused by mutant tau in rTg4510 mice (Ramsden et al., 2005; Santacruz et al., 2005). Indeed, DOX administration, which suppresses tau P301L expression (Figure 3), also reduced the fraction of degenerating dendrites (Figure 4A; log-rank test, p<0.01, n = 43 dendrites from 20 animals in DOX group). DOX administration also slowed the degeneration of axons, although losses did reach untreated rTg4510 levels by the end of our experimental recording period at 55 weeks old (Figure 4B; log-rank test, p<0.05, n = 49 axons from 20 animals in DOX group). As such, by longitudinally tracking neurites across the development of pathology, we observed the dynamics of the neurodegenerative process at the level of individual dendrites and axons.

**Figure 4.**
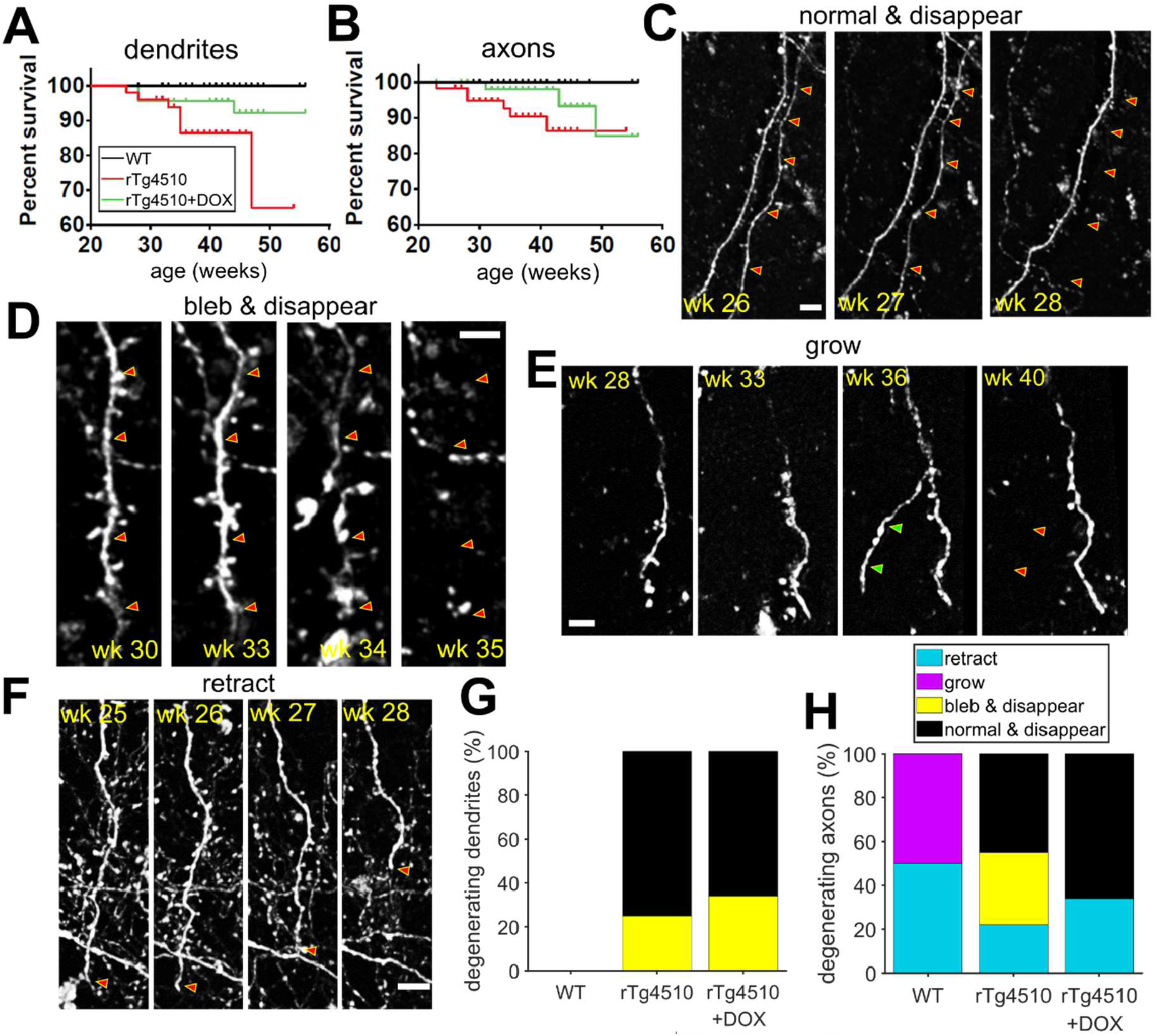
Tauopathy-induced degeneration of dendrites and axons. **(A)** Survival curve for all imaged dendrites in WT, rTg4510 and DOX-treated rTg4510 animals. Ticks indicate data censored due to limitations of imaging rather than neurodegeneration. **(B)** Survival curve for axons in same animals as in (A). **(C)** Example image sequence subset showing disappearance of a dendrite which had healthy morphology one week before (approximate location noted by red arrowheads in all images). The animal’s age at the time of image acquisition is noted for each image. Note the continued presence of a neighbouring dendrite. All image scale bars, 5 μm. **(D)** Image sequence showing a dendrite undergoing profound blebbing (week 34) ahead of disappearance (week 35). **(E)** Image sequence showing axon undergoing growth (wk 36-green arrowheads) and then retraction (week 40-red arrowheads) of a new branch. **(F)** Image sequence showing retraction of an axon over successive weeks (axon tip noted by red arrowhead). **(G)** Stacked bar chart showing the percentage of degenerating dendrites in each morphological category for each group of animals. There were no degenerating dendrites in WT animals. **(H)** Percentages of axons in each group of animals showing different forms of morphological plasticity. In addition to disappearance, some axons in WT animals showed retraction and growth.

Degeneration of neurites occurred heterogeneously, with a variety of morphological changes occurring ahead of neurite disappearance. The majority of degenerating dendrites disappeared between adjacent imaging sessions (1 week interval), having had seemingly healthy structure the week before (normal & disappear; Figure 4C). The other dendrites that were lost had a classically unhealthy appearance, showing increasingly blebbed and/or beaded morphology 1-3 weeks preceding their disappearance (bleb & disappear; Figure 4D). Although the total number of degenerating dendrites was reduced in the DOX-treated mice (Figure 4A), the proportion of morphologies prior to disappearance was similar to that in untreated rTg4510 mice (Figure 4G). By contrast, in WT animals, dendritic backbone structure was extremely stable with no dendrites being lost (Figures 4A&G).

Axons in WT animals did show some plasticity of backbone structure in the form of occasional branch retraction (Figure 4E) and growth (Figure 4F). However, across the WT animals, this remodelling was balanced between retraction and growth, and occurred without morphological signs of dystrophy (Figure 4H). In contrast, we observed no new growth of axons in rTg4510 animals, but there was still clear loss of axons (Figure 4H). These axonal losses were driven by a mixture of retraction and complete disappearance (Figure 4H). As with dendrites, some axons showed dystrophy gradually, with increasingly blebbed morphology in the weeks before disappearance, while others showed no signs of pathology before disappearing between imaging sessions only 1 week apart (Figure 4H). DOX-treated rTg4510 animals similarly had no new growth of axons and the losses were a mixture of retractions and rapid disappearance (Figure 4H). We did not observe blebbing axons in DOX-treated rTg4510 animals in apparent contrast to untreated rTg4510s, but numbers were too small to be certain of this difference.

The neurite losses in rTg4510s occurred at widely varying times. Indeed, it is notable that, while some neurites degenerated quite early in disease progression (<25 weeks old), the disappearance of others is scattered in time through to the end of our imaging period (55 weeks old) (Figure 4A&B). Furthermore, most of the imaged neurites survived to the end of the imaging with little sign of dystrophy, even when they were neighbouring others that had undergone degeneration (Figure 4C). This highlights the variability of the degeneration at the neuronal level and the fact that individual neurites and/or cells are selectively vulnerable.

### Increased loss of spines just prior to dendrite loss

Synaptic dysfunction has been implicated in neurodegenerative disease in rTg4510 mice (Jackson et al., 2017; Kopeikina et al., 2013) and more widely (Selkoe, 2002). Furthermore, some mechanisms that can promote neurite degeneration have been linked to synaptic activity and function (Miyamoto et al., 2017). To investigate potential links between synaptic characteristics and neurite degeneration, we took advantage of our longitudinal approach to align degenerating neurites in time to the week of their disappearance. This allowed us to ask what was happening to synaptic structures in the weeks preceding degeneration. Since each degenerating neurite had at least one other surviving neurite nearby, we also aligned these non-dystrophic neurites in time to directly compare the differences between synaptic characteristics in degenerating and non-degenerating neurites. To measure gains and losses of synaptic structures in dendrites, we tracked each dendritic spine across all the imaging sessions. Because individual neurite degeneration occurred at varying stages of the experiment, there were a variable number of imaging weeks preceding each degeneration event. Therefore, we fitted GAMMs to the data to assess synaptic effects across the populations of degenerating neurites, and matched non-degenerating comparators. Spine turnover dynamics in dendrites destined to be lost were well-matched to those in non-degenerating dendrites in the periods well before the moment of degeneration (Figure 5A; GAMMs based on 20 axons from 9 animals). However, we found that there was an increase in turnover of spines that began just before (∼2 weeks) dendritic degeneration (Figure 5A). This contrasts with non-degenerating dendrites, which maintained stable spine turnover levels through this period. The increase in turnover was driven exclusively by a large increase in the number of spines being lost in the period just preceding dendrite loss (Figure 5B). Indeed, several dendrites lost close to 100% of their existing spines in the week or two before their disappearance. In contrast, the rate of addition of new spines (gains) was maintained during this period (Figure 5C). To assess the effect of this imbalance in gains and losses on spine numbers, we measured the week-to-week changes in spine density (Figure 5D). These data show that there is an acute and rapid loss of synaptic structures in dendrites that immediately precedes death of that dendrite (Figure 5D; repeated measures ANOVA: F(11,55)=4.01, p=0.001 for time-degeneration interaction; post-hoc Tukey: week -1 non-degenerating vs degenerating, p = 0.03).

**Figure 5.**
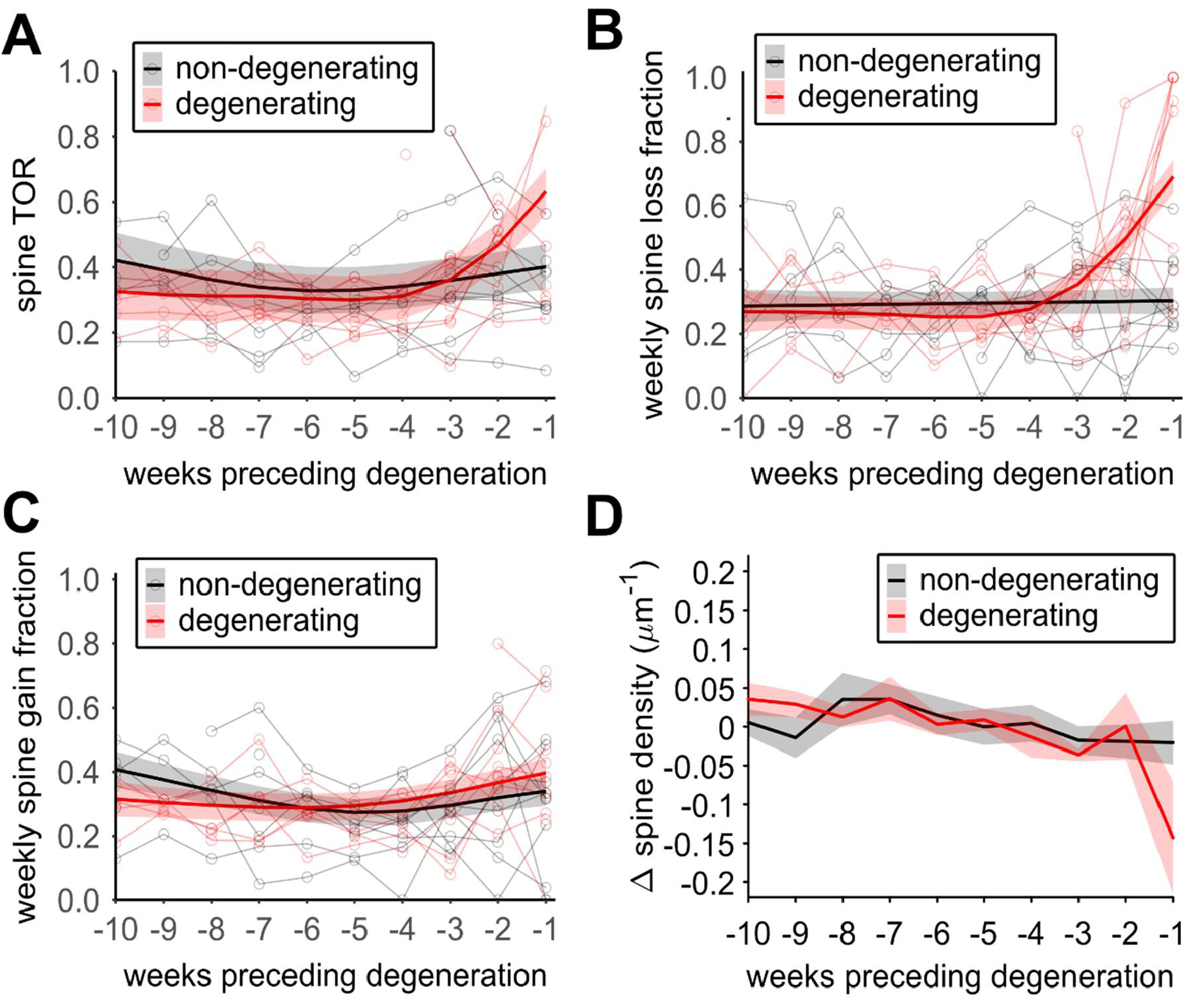
Loss of spines just preceding dendritic degeneration. **(A)** Weekly turnover ratio of dendritic spines for individual branches leading up their degeneration (red, open symbols, n=10 dendrites from 9 animals) alongside equivalent turnover of non-degenerating dendrites (black, open symbols, n=10 dendrites from 9 animals). Each degenerating dendrite has an equivalent non-degenerating dednrites from the same field of view. Overall effects of degeneration are modelled in a GAMM (solid lines, shaded error is 95% confidence intervals). **(B)** Weekly losses of spines ahead of degeneration for same dendrites as (A). **(C)** Weekly gains of new spines ahead of degeneration for same dendrites as (A). **(D)** Week-by-week changes in spine density leading up to the point of degeneration compared to non-degenerating neighbour dendrites (mean+/-SEM).

### Presynaptic bouton turnover is suppressed progressively ahead of axon degeneration

Next, in a similar way, we assessed structural plasticity of TBs in degenerating and matched non-degenerating axons (Figure 6). We found strikingly different effects in axons compared to dendrites. Whereas in dendrites, the spine dynamics were increased in the week or two before degeneration (Figure 5A), the rate of presynaptic bouton turnover starts to decrease ∼2 months ahead of axon loss, maintaining a steady decline up to the point of degeneration (Figure 6A; GAMM based on 18 axons from 7 animals). Turnover rates in non-degenerating axons is relatively stable during the same period (Figure 6A). This reduced TB turnover in degenerating axons is driven by a decrease in both loss (Figure 6B) and addition (Figure 6C) of boutons. This suggests that presynaptic structures display chronic over-stabilisation in axons that go on to degenerate. Since there is a decrease in the rate of emergence of new TBs and the loss of existing ones, it is perhaps not surprising that there were no clear week-by-week changes in the TB density leading up to degeneration (Figure 6D; repeated measures ANOVA: F(11,22)=1.01, p=0.47). As such, it appears that, in contrast to dendrites that undergo dramatic synaptic losses just ahead of degeneration, axons are predominantly characterised by a progressive denigration of structural plasticity that starts many weeks before they die.

**Figure 6.**
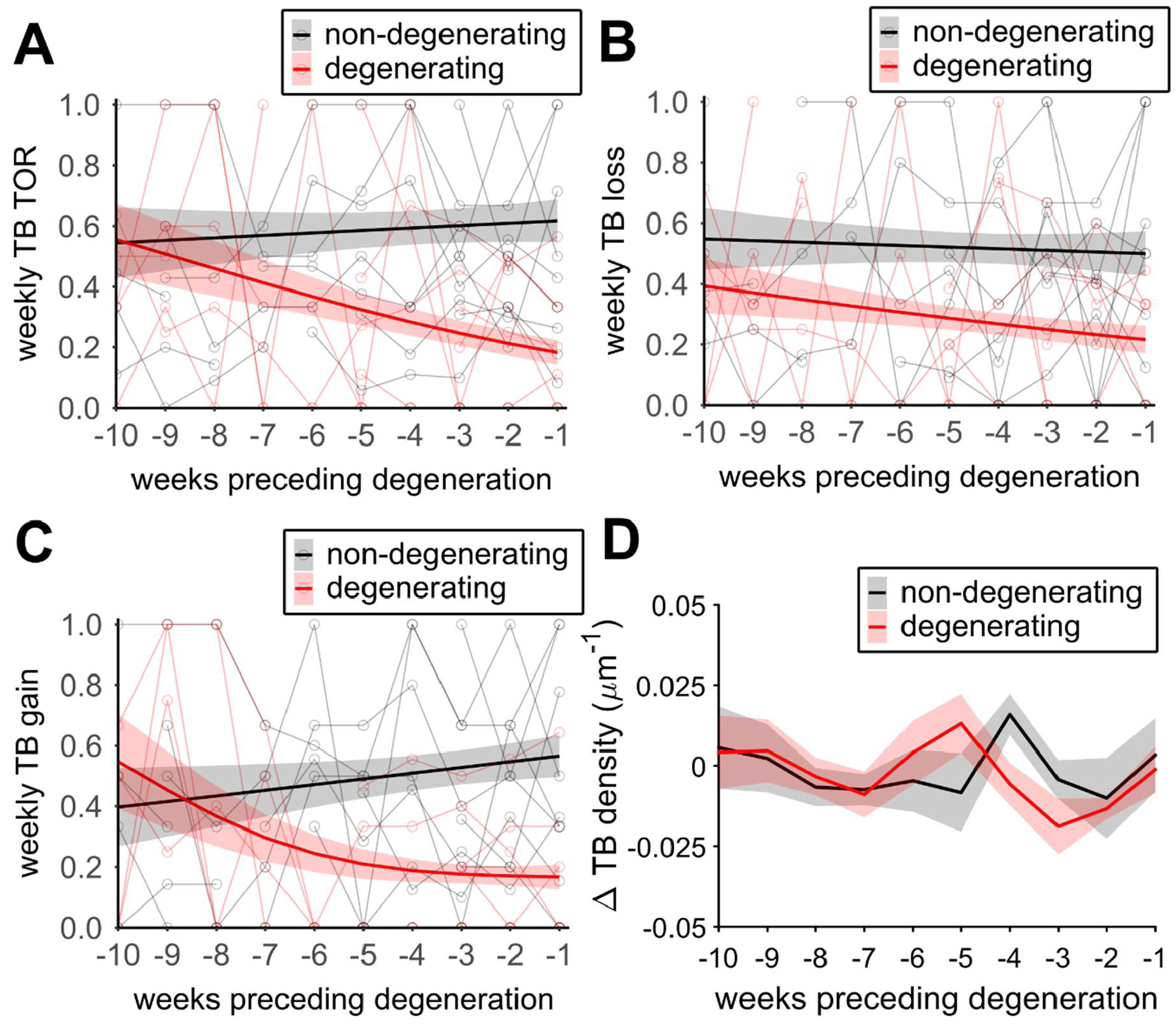
Progressively reduced turnover of presynaptic TBs in the months preceding axonal degeneration. **(A)** Weekly turnover ratio of TBs for individual branches leading up their degeneration (red, open symbols, n=9 axons from 7 animals) alongside equivalent turnover of non-degenerating dendrites (black, open symbols, n=9 axons from 7 animals). Each degenerating axon has an equivalent non-degenerating axon from the same field of view. Overall effects of degeneration are modelled in a GAMM (solid lines, shaded error is 95% confidence intervals). **(B)** Weekly losses of TBs ahead of degeneration for same axons as (A). **(C)** Weekly gains of new TBs ahead of degeneration for same axons as (A). **(D)** Week-by-week changes in TB density leading up to the point of degeneration compared to non-degenerating neighbour dendrites (mean+/-SEM).

## Discussion

By repeatedly imaging large numbers of axons and dendrites each week over months, we have shown that tauopathy-driven degeneration of the parent axon or dendrite is associated with aberrant turnover of synapses in the weeks leading up to its death (Figures 4&5). Our longitudinal approach to imaging of individual axons and dendrites shows that there is a huge diversity of effects of tauopathy at varying times in different cells. Overall, as predicted from previous cross-sectional measurements of synapse density in rTg4510 animals (Crimins et al., 2012; Jackson et al., 2017; Kopeikina et al., 2013), we found that the density of both presynaptic TB and postsynaptic dendritic spines is decreased as the pathology progresses (Figures 1&2). These decreases in overall synapse density are driven partly, but not exclusively, by degeneration of dendrites and axons is distributed widely across the time-course of disease. Despite the widely varying times of degeneration, our tracking of individual neurites allowed us to retrospectively measure their properties as they approached their death and compare them to healthy neighouring cells. We found that neurite degeneration is preceded by aberrant structural synaptic plasticity in a cell-specific way that is markedly different in dendrites and axons.

In many degenerating dendrites, there is a dramatic loss of dendritic spines in the week or two before they disappear (Figure 5). This suggests that there is a rather rapid final decline in dendritic functional integrity just before tau-driven degeneration. We did observe some classic signs of pathology, such as swelling or blebbing, in a minority of degenerating dendrites, but these signs were only ever manifested in the 2-3 weeks before dendrite loss (Figure 4). Indeed, most dendrites that were lost did not show these classic pathological symptoms before degeneration. It is possible that the timing of our imaging means that we missed the emergence of pathology. However, our weekly imaging interval means that, if we did miss blebbing or swelling, it must have only occurred in the a few days before dendritic destruction. Retraction, growth or partial loss of dendrites was extremely rare in rTg4510s and was never observed in WT animals, suggesting that the entire dendrite is lost at once. These rapid and complete losses of a specific dendrite within a field of view could well be linked to death of the parent neuron. A similar dendritic degeneration phenomenon has been observed in the 3xTg-AD mouse line, which expresses amyloid and tau pathological mechanisms (Bittner et al., 2010). In that study, cortical neuronal loss, and associated dendritic degeneration, occurred early in pathological progression, prior to detection of plaques or NFTs (Bittner et al., 2010). However, both diffuse amyloid and phosphorylated tau have been detected at these early stages in the cortex of the 3xTg-AD mouse (Cai et al., 2012), which leaves open the possibility that either or both drive the observed neuronal loss. Notably, many mouse models of amyloidosis do not exhibit cell death or neurite loss despite overt amyloid pathology, leaving tau more directly implicated in neuronal cell death (Lee and Han, 2013; Stephen et al., 2019). Intriguingly, some of the dendritic losses we observed in rTg4510s also occurred in relatively young animals, at a stage that precedes the major emergence of NFTs. Taken together, this would suggest that tau-based pathology can drive dendritic (and potentially, neuronal) degeneration independent of insoluble NFTs.

Axons also showed tau-dependent degeneration, with losses occurring across wide-ranging times, starting as early as 23 weeks old (Figure 4). Overall, more dendrites were lost than axons, aligning with the more severe dendritic synapse losses here (Figures 1 & 2) and previously observed (Jackson et al., 2017). However, as with dendrites, most of these axonal losses occurred without any overt signs of dystrophy in preceding weeks, although a minority did display blebbing ahead of degeneration (Figure 4). However, in contrast to dendrites, some axons did display plasticity of their backbone structure. Even in WT animals, we observed a small and balanced number of axonal retraction and growth events. In transgenic animals, no growth was observed but some axons did appear to retract. Even so, ∼60% of axons that disappeared seemed to have normal structure in the preceding imaging session, just one week before (Figure 4). This aligns with the idea that a majority of tauopathy-induced neurite disappearance is associated with acute loss of large sections of neurite, or perhaps the entire cell. Alternatively, however, the disappearance of axons from a field of view could be due to a relatively rapid “dying back” mechanism, which has been previously associated with neurodegenerative disease (Adalbert and Coleman, 2013; Kneynsberg et al., 2017), if the degenerating axons died back sufficiently rapidly that it occurred within the week-long period between imaging sessions. We detected relatively few “dying back” axons but, if this process is indeed rapid, then we may have needed much larger fields of view or more frequent imaging sessions to detect them.

The cell-autonomous nature of both dendritic and axonal degeneration is highlighted by the fact that each degenerating neurite was near other imaged neurites which showed no signs of pathology. Indeed, most of the imaged neurites in the rTg4510 animals survived the entire imaging time-course without degenerating. The low rates of detection of degenerating neurites (∼1% per week) may seem counter-intuitive given the overt loss of cortical volume overall in rTg4510s (Santacruz et al., 2005)(Figure 3B), but these degeneration events are, of course, cumulative and therefore could add up to a substantial effect over time.

Reduction of tau expression has been suggested as potential therapeutic strategy for tauopathy-driven neurodegeneration. We tested the link between continued tau expression and neurodegeneration by dosing the rTg4510 animals with DOX starting at 18, 22, 26 and 30 weeks of age. DOX reduced expression of mutant tau by ∼50%, similarly in all treatment groups (Figure 4A). In line with other studies (Holmes et al., 2016; Wang et al., 2018), we found that the effect of reducing tau P301L expression on subsequent appearance of classical markers of tau pathology and neurodegeneration was dependent on when DOX administration was started. The earliest administration, at 18 weeks, reduced PG5 accumulation, ameliorated cortical atrophy and partially rescued tau-induced reductions in synapse density (Figure 3B-G). However, there was no discernible beneficial effect if DOX was started after 22 weeks (Figure 3E). This confirms that there is a stage, relatively early in tauopathy, beyond which the progression of pathology will continue unabated even if the addition of aberrant tau is curtailed (Helboe et al., 2017). In line with the partial rescue of classical markers of pathology, DOX administration slowed, but did not stop, degeneration of individual axons and dendrites (Figure 4A). A recent study has shown that there is disruption of off-target genes in the rTg4510 animals at the site of transgene insertion (Gamache et al., 2019). It was suggested that this genetic disruption may drive rTg4510 pathology independent of tau (Gamache et al., 2019). The fact that DOX (and consequent reduction of tau P301L expression) did have an ameliorative effect at young ages in our study suggests that tau expression must be contributing to pathological progression at that stage (Figure 3&4). This aligns with previous studies that showed suppression of tau expression in rTg4510 animals from conception to various stages of adulthood, which effectively eliminates tau P301L effects but leaves other genetic disruption intact in the responder line, completely eliminates pathological progression until DOX is removed (Han et al., 2012; Yang, 2013; Hunsberger et al., 2014; Helboe et al., 2017). This suggests that tau P301L is the predominant driver of pathology. However, the lack of an available genetically-matched control leaves open the possibility in our experiments, and those of others (Helboe et al., 2017; Holmes et al., 2016; Wang et al., 2018), that the lack of DOX effect when administered later is because genetic factors other than tau contribute to later rTg4510 pathology.

Abnormal synapse function is a well-characterised phenomenon associated with tauopathy and other forms of neurodegenerative disease. Our longitudinal imaging allowed us to capture structural synaptic plasticity in axons and dendrites in the weeks leading up to their degeneration. This analysis revealed that degeneration is preceded by neurite-specific changes in synapse plasticity. In dendrites, a reduction in synapse density in the 2 weeks ahead of degeneration is driven by a dramatic increase in the removal of spines. Loss of postsynaptic input may be an early symptom of the cellular pathology leading up to neuron death or may itself be a driver of degeneration. Indeed, synaptic weakening, and particularly long-term depression, have been associated with the development of pathology in several models of neurodegenerative disease, including rTg4510s (Crimins et al., 2012; Eckermann et al., 2007; Jackson et al., 2017; Thies and Mandelkow, 2007; Yoshiyama et al., 2007). Our data suggest that at least some of this synapse depression is very closely linked in time to dendrite degeneration. This short time window should shape the design of any intervention aimed at ameliorating degenerative pathology by normalising synaptic strength. Such an intervention should probably be targeted at halting the onset of synaptic depression rather than reversing it. Interestingly, even during the period of intense reduction in synapse density, new spines were still being added at a normal rate. This suggests that the mechanisms for creating new synapses are still functional as dendrites enter the final stages of cellular pathology.

Perhaps the most striking result from our studies is the fact that degeneration-associated synaptic plasticity in axons is very different from that in dendrites. There is a slowly progressive reduction in structural synaptic plasticity that emerges from approximately two months ahead of axon loss (Figure 6A). In stark contrast to the increased loss of dendritic spines, axons destined for degeneration actually undergo a reduced rate of synapse loss (Figure 6B). These reduced losses do not, however, drive a dramatic increase in TB density because, along the same time-course, the addition of new TBs is also inhibited (Figure 6D). Overall, this may contribute to a slow attrition of presynaptic density because gains appear to be slighter more inhibited than losses. In fact, the gained fraction of axonal synapses drops to almost zero as the point of degeneration approaches. This suggests a scenario in which presynaptic sites become abnormally stable ahead of degeneration. The relative lack of plasticity may reflect an inability of these cells to participate in the normal changes in neuronal connectivity that underlies ongoing brain function. Since turnover of synapses has been related to learning and memory formation, it is feasible that pathological synaptic overstabilisation in axons could relate to cognitive symptoms associated with disease progression.

Our findings give weight to the idea that aberrant tau may exert pathological influence on synapses via different mechanisms is pre- and postsynaptic compartments. Under healthy physiological conditions, tau acts as microtubule-stabiliser within the axon (Goedert and Spillantini, 2011). Hyperphosphorylation of tau leads to dissociation from the microtubules, decreasing their stability. Microtubule instability has been shown to negatively impact axon structure and fast axonal transport of cellular organelles, which could impact presynaptic structural plasticity. Furthermore, direct interaction of hyperphosphorylated tau with presynaptic vesicles was shown to mediate aberrant synaptic function (Zhou et al., 2017), which could impact plasticity. It is possible that over-stabilisation of presynaptic sites plays a role in driving subsequent axon degeneration. The relatively prolonged period of aberrant plasticity before degeneration opens a potential window of opportunity to test whether rescue of plasticity defects may have ameliorative benefits for axons. In contrast to axons, tau is not usually found within dendrites, but it is mis-targeted there following its hyperphosphorylation (Zempel and Mandelkow, 2014). This pathological tau is thought to mediate postsynaptic abnormalities by modulating Fyn kinase activity that impacts NMDA receptors (Ittner et al., 2010). Given the important role of NMDA receptors in triggering various forms of synaptic plasticity as well as cell death pathways (Amadoro et al., 2006; Hardingham and Bading, 2010), it is tempting to suggest that this mechanism may be involved in the degeneration-associated changes in dendrite plasticity described here.

We anticipate that there are synaptic connections between the cortical cells within the population that were labelled in this study, which raises the question of how such different plasticity dynamics on either side of the synapse can be reconciled. Opposite shifts in dynamics of synapse turnover could represent compensatory mechanisms for what is happening on the other side of the synapse, or perhaps in response to sudden loss of synaptic partners due to cell death. It will be important to image locations where labelled axons and dendrites make synapses with each other to directly assess the interplay between the differential pathological dynamics of pre- and postsynaptic compartments as they undergo degeneration.

## Methods

### Animals

The rTg4510 mice line expresses a reversible transgene expressing the 4R0N tau isoform with the P301L mutation under control of doxycycline (DOX)-dependent promoter (Santacruz et al., 2005). Verification of genotypes were assessed by a standardized PCR assay for activator and responder transgenes (Santacruz et al., 2005). We compared groups of animals; wild-type (WT) littermates, rTg4510 animals given vehicle and rTg4510 animals given DOX. Animals were randomly allocated to four batches in which imaging was started at differing ages (20, 24, 28 and 32 weeks). There were 5 animals in each group within each batch, for a total of 60 animals. To turn off transgene expression, doxycycline was administered through an initial oral bolus (10mg/kg) following cranial window surgery; mice were then fed a doxycycline-containing diet (200 mg/kg of dietary chow) throughout the imaging period. All mice were given ad libitum access to food and water and maintained in a 12-hour light-dark cycle. All procedures were conducted by researchers holding a UK personal licence and conducted in accordance with the UK Animals (Scientific Procedures) Act 1986, and subject to internal ethical review. All experiments and analyses were completed blind to genotype or treatment group.

### Surgery

Two surgeries were performed on each animal. In the first surgery, performed when animals were aged between 15-17 weeks under 1.5% (v/v) isoflurane induced anaesthesia, an adeno-associated virus (AAV) serotype 2 expressing GFP (10^10^ GU; Vector Biolabs) was injected into layer 2/3 (300µm below the dura) to enable visualisation of cortical neurons. Viral injections of 0.3µl per site, at three sites running approximately 1mm from and parallel to the midline were performed. Mice were given pre-operative intra-peritoneal injections of dexamethasone (30 mg/kg) to reduce brain swelling and buprenorphine (5 mg/kg) for analgesia. Performing these viral infections at an early, well-defined pre-degenerative age ensures that similar population of neurons is infected in all animals. Mice were allowed to recover for at least 2 weeks before the second surgery in which a cranial window was implanted to allow *in vivo* imaging. The cranial window was implanted 2 weeks ahead of the first imaging session. Therefore, this surgery was performed at different ages for the four different batches of animals (18, 22, 26 and 30 weeks). Alignment of window implantation with imaging age maximised the utility of the cranial window for the target imaging ages in each batch. Mice were again given pre-operative intra-peritoneal injections of dexamethasone (30 mg/kg) to reduce brain swelling and buprenorphine (5 mg/kg) for analgesia. For window implantation, under 1.5% (v/v) isoflurane induced anaesthesia, the skull was exposed and a 4-5mm diameter section overlying the somatosensory cortex was removed (AP +1.4, ML -3.0). A 5mm glass coverslip was placed over the craniotomy and sealed with glue and dental cement containing gentamicin. A metal screw was implanted on the contralateral skull to aid window stability. Further dental cement was then added over the remaining exposed skull, and a stainless-steel bar (<500 mg) implanted on to the anterior contralateral window to enable accurate positioning on the two-photon microscope. Mice were left to recover for at least 2 weeks prior to the first imaging session.

### Longitudinal imaging

Longitudinal two-photon imaging was performed as previously described (Jackson et al., 2017). Animals were imaged weekly starting at 20, 24, 28 and 32 weeks for each batch. Staggering the onset of imaging allowed coverage of ages prior to and throughout widespread cortical pathology. A two-photon microscope (Prairie Technologies) equipped with a tuneable Ti:Sapphire pulsed laser (∼100 fs pulsewidth, MaiTai HP, SpectraPhysics) and PrairieView acquisition software was used for all imaging experiments. For each imaging session mice were positioned and secured on the microscope stage via the implanted steel head bar. Mice were maintained under anaesthesia throughout imaging with 3-5% isoflurane. The body temperature was measured and maintained above 35°C using a rectal probe and heating blanket. Lacri-lube was added to the eyes to stop them from drying out. A 10x objective (NA=0.3, Olympus) was used to identify superficial blood vessels as fiducial markers, enabling the relocation of regions of interest (ROIs). A 40x water immersion objective (LUMPlanFI/IR, NA=0.8, Olympus) was then used to acquire Z-stacks of each ROI (75 µm x 75 µm, 512 x 512 pixels, step size = 0.5 µm) per animal. Excitation power at 910 nm was kept below 35mW at the sample to avoid phototoxicity. GFP emission was collected through a 525/25nm filter. Initially 5-10 ROIs containing neurites were chosen and images acquired ensuring ROIs were separated by at least 100µm. In each subsequent imaging session, each ROI was relocated and imaged for up to 26 sessions. There were no differences between genotypes in image signal-to-noise ratio, measured by mean:standard deviation of the upper 15^th^ percentile of pixel intensity values. Animals were removed from subsequent imaging sessions if there was significant clouding of the cranial window that made it impossible to re-locate and/or visualise neurites. After the final imaging session, each animal was sacrificed by cardiac perfusion with 4% paraformaldehyde.

### Spine and terminaux bouton structure analysis

*In vivo* two-photon images were converted into stacks with ImageJ (National Institutes of Health) and the registered in XY using the StackReg plugin to account for any movement or drift during imaging (Thevenaz et al., 1998). The stacks were deconvolved with Huygens Deconvolution software (Scientific Volume Imaging) using a quick maximum likelihood estimation with an experimentally defined point-spread-function. Each ROI Z-stack has all sessions aligned in 3D using “Least squares” landmark registration on MIPAV (National Institutes of Health). Aligned ROIs were then converted into a 4D hyper-stack (XYZT) enabling the optimal medium for spine and bouton analysis. Axons and dendrites were distinguished by their morphology, with a straighter, thicker dendritic shaft vs tortuous thinner axonal process as described previously in the literature (De Paola et al., 2006; Majewska et al., 2006). All dendritic spines were counted, while only terminaux boutons were counted on axons. Synaptic components were counted manually following a ruleset from previous literature (Holtmaat and Svoboda, 2009; Jackson et al., 2017) using the Cell Counter plugin on ImageJ. Further data manipulation and analysis was carried out using MATLAB (Mathworks).

### Histology

Perfused-fixed brains were coronally dissected into three segments using an adult mouse brain matrix (slot #5 and #11 AP; RBM-2000C: ASI Instruments, USA). These segments were processed using the Tissue TEK® VIP processor (GMI Inc, USA) and embedded in paraffin wax. The middle segment was used to cut 8 μm serial sections using rotary microtomes (HM 200 and HM 355; Thermo Scientific, Germany) which were mounted on glass slides. Coronal sections representing approximately Bregma -1.50 (AP) were selected for immunohistochemistry using primary antibodies specific for phospho-tau (PG-5, tau phosphorylated at s409; 1:8000, from Peter Davies) and microglia (Iba-1: 1:6000, Wako Chemicals GmbH, Germany). Following de-paraffinisation and rehydration of the tissue, antigen retrieval was performed using the Lab Vision PT module system (Thermo Scientific, UK), where sections were heated to 100°C for 20 min in citrate buffer (TA-250-PM1X; Thermo Scientific, UK). After cooling in dH_2_O, the slides were transferred to the Lab Vision Autostainer 360 (Thermo Scientific, UK) where the following incubations were performed: 10 min in H_2_O_2_ (0.03%); 30 min in normal goat serum (1:20; Vector Laboratories, USA); 60 min in primary antibodies; 30 min in biotinylated goat anti-mouse or anti-rabbit IgG (1:200, PA-920 or BA-1000; Vector Laboratories, USA); 30 min avidin-biotin complex solution (PK-7100; Vector Laboratories, USA); 5 min in 3,3′-diaminobenzidine (SK-4105; Vector Laboratories, USA). Apart from the last two steps, PBS with 0.05% Tween-20 (PBS-T) was used for diluting reagents and washes between steps. Sections were then counterstained with haematoxylin before dehydration and cover-slipping. Stained sections were digitised using the Scanscope XT slide scanner (Aperio, USA) at 20× magnification. Imagescope software (version 11.1.2.760; Aperio, USA) was used to view the digitised tissue sections and delineate boundaries of the hippocampus and overlying cortex (including barrel field of the somatosensory cortex) for both the right (cranial window) and left (contralateral) side of brain. Immunoreactivity for PG-5 positive tau pathology and Iba-1 positive microglia within the regions of interest was quantified using the positive pixel algorithm (Imagescope, version 11.1.2.760; Aperio, CA, USA) and expressed as a percentage of the total area.

### Quantitative reverse transcription PCR

Expression of transgenic tau was analysed by reverse transcription quantitative PCR using post-mortem fixed cortical samples from individual brains as previously described (Blackmore et al., 2017). Data were analysed using the ΔΔCt method using GAPDH as the reference gene and were normalised to a known untreated rTg4510-positive sample from late stage pathology. Of note, one animal genotyped as WT by conventional PCR, showed mildly (27% of rTg4510 standard) elevated tau P301L level by quantitative PCR. However, this animal showed WT levels of PG5 and IBA-1 staining and normal brain weight. Therefore, we assumed an unexplained error in quantitative PCR in this sample, and retained this animal as WT genotype.

### Statistical analysis

Generalized Additive Mixed Models (GAMMs) were used to assess any changes in synapse density or dynamics over time (Figures 1,2,5&6). GAMMs extend typical regression methods to estimate the relationship between a dependent variable (e.g. spine density) and specified predictors (e.g. genotype, age, animal, batch)(van Rij et al., 2019; Shadish et al., 2014; Wood, 2017). This relationship is modelled based on a smooth function rather than a typical linear regression. As such, it allows deviation from a linear relationship between predictors and dependent variables, which is potentially important because effects of neurodegeneration may vary between neurons at different stages of disease. Furthermore, the influence of repeated measures from individual animals and multiple neurites (likely neurons) within each animal, as well as different batches, can be included as random effects. Also, unlike repeated measures ANOVAs, GAMMs allow us to incorporate data from individual subjects at different (not necessarily matching) timepoints. This is important because of the varying time-course of imaging in different animals, the fact that some neurites degenerate (and therefore no longer contribute to population density) and because there were a few occasions when image sessions were lost for technical reasons (e.g. imaging system or anaesthesia complications). GAMMs were fitted using gam function of the *mgcv* package in R (Wood, 2017). Changes in mean synapse density over time were fitted using the general form:

*synapse property ∼ s(age, by group) + group + (1*|*individual_neurite,age) + (1*|*individual_neurite) + (1*|*animal,week) + (1*|*batch,group)*

The smoothing spline (*s*) fitted over *age* individually for each *group* (i.e. genotype or degenerating/not degenerating) with no prespecified number of smoothing knots (since the trajectory across time was not known *a priori*). Variability associated with repeated measures of individual *neurites, animals* and *batch* over *age* were included as random effects. GAMMs for spine properties were fitted to Tweedie distributions to account for occasions when they fell to zero (Supplementary Figure 1A). GAMMs for TB density was fitted to a Gaussian distribution (Supplementary Figure 1C) and TB turnover properties were fit using a beta distribution. Choice of the modelled distribution (and link function) was based on subjective assessment of linearity of QQ-plots and maximising deviance explained by the model (Supplementary Figure 1B&D).

Histological and qPCR data was assessed via ANOVA followed by Tukey post-hoc testing when a significant effect (P<0.05) was found. Neurite survival was analysed by Kaplan-Meier curves and differences between genotypes tested using log-rank tests.

## Author contributions

JDJ, JSJ, MLH, JI, MCA and MJO designed the study; JDJ and JSJ conducted the 2-photon imaging and image analysis; SM and ZA conducted the histology and analysis; TM coordinated and managed transgenic animals; JDJ, JSJ and MCA conducted data analysis; MF advised and conducted statistical analyses. All authors contributed to the manuscript preparation.

## Acknowledgements

This work was funded by Eli Lilly and Co. JDJ was funded by a BBSRC-CASE PhD studentship (1370828). MCA’s laboratory was funded by the Medical Research Council (MR/J013188/1) and EUFP17 Marie Curie Actions (PCIG10-GA-2011-303680).

## Supplementary Material

**Supplementary Figure 1.**
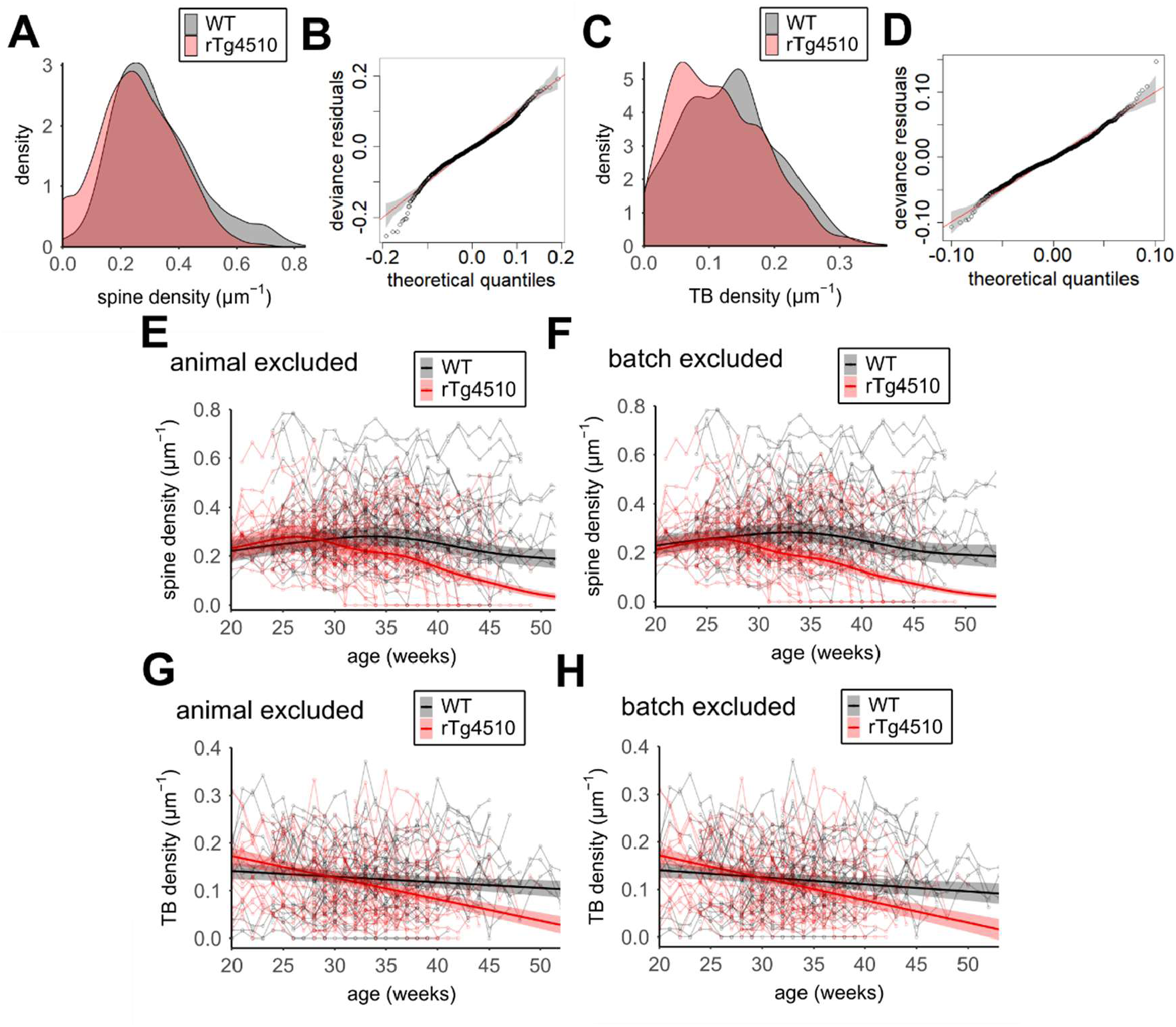
Statistical analysis based using GAMMs. **(A)** Density distribution of spine density values from all dendrites and all imaging timepoints in WT and rTg4510 animals. **(B)** QQ plot of residuals from spine density GAMM modelled on Tweedie distribution. **(C)** Density distribution of axonal TB density values across all imaging timepoints in WT and rTg4510 animals. **(D)** QQ plot of residuals from TB density GAMM modelled on gaussian distribution. **(E)** GAMM predictions for changes in spine density (thick line with shaded 95% confidence limits) in the absence of *animal* as a random variable. Note there is a significant but small deviation in predictions for each genotype from the full model (Figure 1E). **(F)** GAMM predictions for changes in spine density (thick line with shaded 95% confidence limits) in the absence of *batch* as a random variable. The exclusion of batch from the model makes almost no difference to predictions for each genotype compared to the full model (Figure 1E)_ **(G)** GAMM predictions for changes in spine density (thick line with shaded 95% confidence limits) in the absence of *animal* as a random variable. Note there is a significant but small deviation in predictions for each genotype from the full model (Figure 1F). **(H)** GAMM predictions for changes in spine density (thick line with shaded 95% confidence limits) in the absence of *batch* as a random variable. The exclusion of batch from the model makes almost no difference to predictions for each genotype compared to the full model (Figure 1F).

**Supplementary Figure 2.**
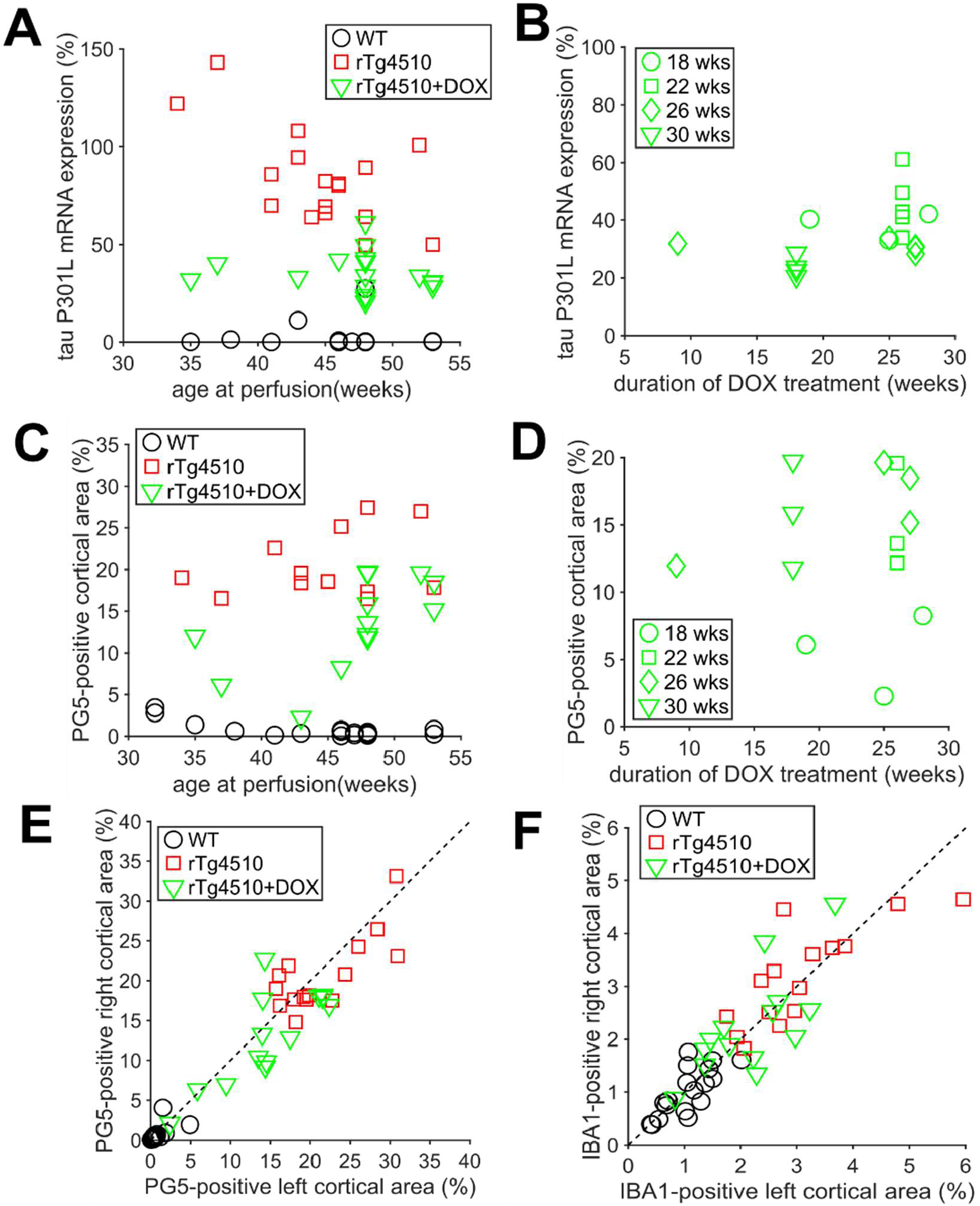
Effects of tau P301L transgene expression level on histopathology. **(A)** Postmortem expression level of tau P301L (qPCR relative to independent standard rTg4510 sample) for individual WT, untreated rTg4510 and DOX-treated rTg4510 animals perfused at varying ages. DOX partially suppresses tau P301L expression similarly across the ages tested, (n(animals)= 16 WT,17 rTg4510,17 rTg4510+DOX). **(B)** Suppression of tau P301L expression in DOX-treated rTg4510 animals is largely independent of the duration of treatment or the age at the start of treatment, (n(animals) = 3,5,5,4 for 18,22,26,30 week groups). **(C)** Histopathological measurement of PG5 levels in individual animals at age of perfusion. **(D)** Duration of DOX treatment appears to have limited influence on PG5 staining. Note that early onset of treatment (20 week group) has the lowest PG5 levels independent of treatment duration. **(E)** PG5 levels are similar in somatosensory cortex in both hemipsheres of all individual animals. **(F)** Coverage of activated microglia, measured by IBA1 staining, is similar in somatosensory cortex in both hemipsheres of all individual animals. The cranial window was always implanted over right somatosensory cortex. For PG5 and IBA1 data, n (animals)=18 (4, 4, 5, 5) WT, 14 (4,3,4,4) rTg4510, 13 (3, 3, 4,3) rTg4510+DOX (animals in each batch started at 18,22,26,30 weeks of age shown in parentheses).

**Supplementary Figure 3.**
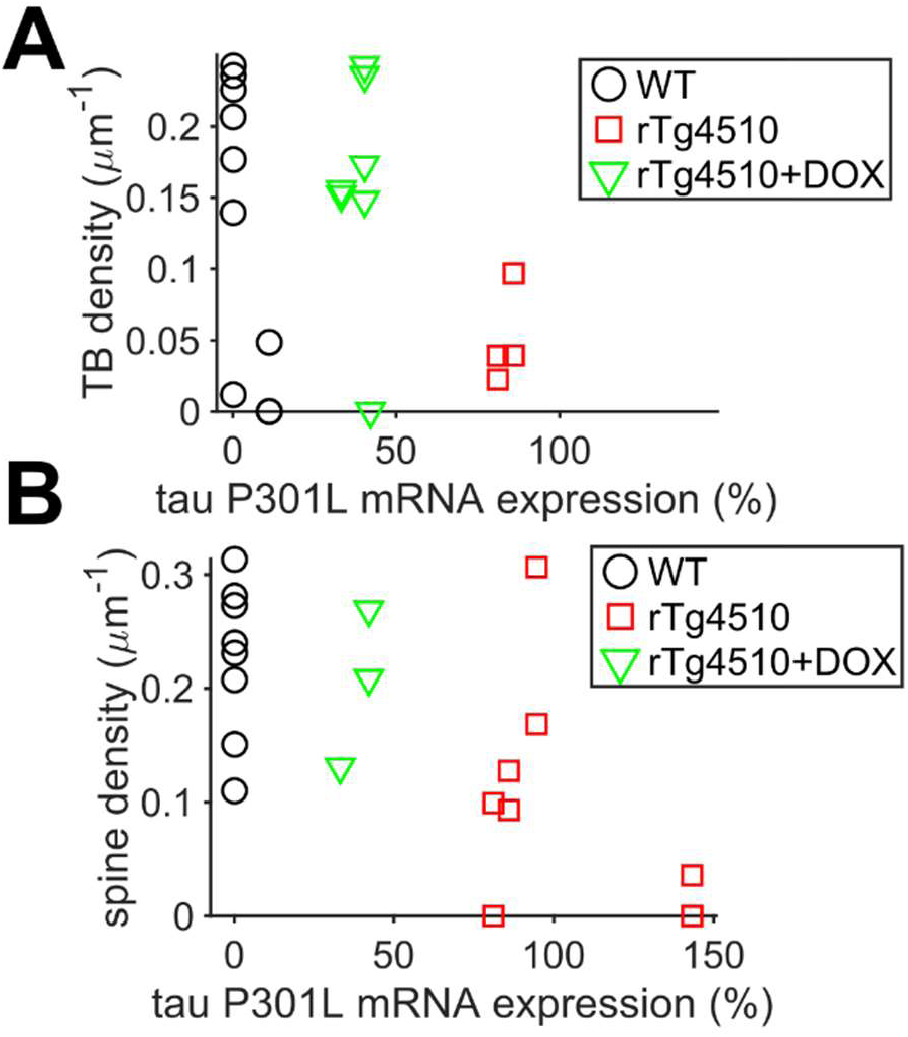
Relationship between tau P301L transgene expression level and synapse density in neurites from WT, rTg4510 and DOX-treated rTg4510 animals. **(A)** Overall, median presynaptic TB density tends to be reduced with increasing tau P301L expression but can vary widely between individual axons even when postmortem expression levels of tau P301L are similar. **(B)** Similarly to TBs, there is substantial variability in the median spine density on individual dendrites from animals with similar postmortem tau P301L expression levels.

## References

Adalbert, R., and Coleman, M.P. (2013). Review: Axon pathology in age-related neurodegenerative disorders. Neuropathol. Appl. Neurobiol. 39, 90–108.

Amadoro, G., Ciotti, M.T., Costanzi, M., Cestari, V., Calissano, P., and Canu, N. (2006). NMDA receptor mediates tau-induced neurotoxicity by calpain and ERK/MAPK activation. Proc. Natl. Acad. Sci. 103, 2898–2897.

Bittner, T., Fuhrmann, M., Burgold, S., Ochs, S.M., Hoffmann, N., Mitteregger, G., Kretzschmar, H., LaFerla, F.M., and Herms, J. (2010). Multiple Events Lead to Dendritic Spine Loss in Triple Transgenic Alzheimer’s Disease Mice. PLoS ONE 5, e15477.

Blackmore, T., Meftah, S., Murray, T.K., Craig, P.J., Blockeel, A., Phillips, K., Eastwood, B., O’Neill, M.J., Marston, H., Ahmed, Z., et al. (2017). Tracking progressive pathological and functional decline in the rTg4510 mouse model of tauopathy. Alzheimers Res. Ther. 9, 77.

Booth, C.A., Witton, J., Nowacki, J., Tsaneva-Atanasova, K., Jones, M.W., Randall, A.D., and Brown, J.T. (2016). Altered Intrinsic Pyramidal Neuron Properties and Pathway-Specific Synaptic Dysfunction Underlie Aberrant Hippocampal Network Function in a Mouse Model of Tauopathy. J. Neurosci. 36, 350–363.

Cai, Y., Zhang, X.-M., Macklin, L.N., Cai, H., Luo, X.-G., Oddo, S., LaFerla, F.M., Struble, R.G., Rose, G.M., Patrylo, P.R., et al. (2012). BACE1 Elevation is Involved in Amyloid Plaque Development in the Triple Transgenic Model of Alzheimer’s Disease: Differential Aβ Antibody Labeling of Early-Onset Axon Terminal Pathology. Neurotox. Res. 21, 160–174.

Crimins, J.L., Rocher, A.B., and Luebke, J.I. (2012). Electrophysiological changes precede morphological changes to frontal cortical pyramidal neurons in the rTg4510 mouse model of progressive tauopathy. Acta Neuropathol. (Berl.) 124, 777–795.

Eckermann, K., Mocanu, M.-M., Khlistunova, I., Biernat, J., Nissen, A., Hofmann, A., Schönig, K., Bujard, H., Haemisch, A., Mandelkow, E., et al. (2007). The β-Propensity of Tau Determines Aggregation and Synaptic Loss in Inducible Mouse Models of Tauopathy. J. Biol. Chem. 282, 31755–31765.

Forner, S., Baglietto-Vargas, D., Martini, A.C., Trujillo-Estrada, L., and LaFerla, F.M. (2017). Synaptic Impairment in Alzheimer’s Disease: A Dysregulated Symphony. Trends Neurosci. 40, 347–357.

Gamache, J., Benzow, K., Forster, C., Kemper, L., Hlynialuk, C., Furrow, E., Ashe, K.H., and Koob, M.D. (2019). Factors other than hTau overexpression that contribute to tauopathy-like phenotype in rTg4510 mice. Nat. Commun. 10, 2479.

Gendron, T.F., and Petrucelli, L. (2009). The role of tau in neurodegeneration. Mol. Neurodegener. 4, 13.

Goedert, M., and Spillantini, M.G. (2011). Pathogenesis of the tauopathies. J. Mol. Neurosci. MN 45, 425–431.

Grutzendler, J., Kasthuri, N., and Gan, W.-B. (2002). Long-term dendritic spine stability in the adult cortex. Nature 420, 812–816.

Han, H.J., Allen, C.C., Buchovecky, C.M., Yetman, M.J., Born, H.A., Marin, M.A., Rodgers, S.P., Song, B.J., Lu, H.-C., Justice, M.J., et al. (2012). Strain Background Influences Neurotoxicity and Behavioral Abnormalities in Mice Expressing the Tetracycline Transactivator. J. Neurosci. 32, 10574–10586.

Hardingham, G.E., and Bading, H. (2010). Synaptic versus extrasynaptic NMDA receptor signalling: implications for neurodegenerative disorders. Nat. Rev. Neurosci. 11, 682–696.

Harris, J.A., Koyama, A., Maeda, S., Ho, K., Devidze, N., Dubal, D.B., Yu, G.-Q., Masliah, E., and Mucke, L. (2012). Human P301L-Mutant Tau Expression in Mouse Entorhinal-Hippocampal Network Causes Tau Aggregation and Presynaptic Pathology but No Cognitive Deficits. PLOS ONE 7, e45881.

Helboe, L., Egebjerg, J., Barkholt, P., and Volbracht, C. (2017). Early depletion of CA1 neurons and late neurodegeneration in a mouse tauopathy model. Brain Res. 1665, 22–35.

Herms, J., and Dorostkar, M.M. (2016). Dendritic Spine Pathology in Neurodegenerative Diseases. Annu. Rev. Pathol. Mech. Dis. 11, 221–250.

Holmes, H.E., Colgan, N., Ismail, O., Ma, D., Powell, N.M., O’Callaghan, J.M., Harrison, I.F., Johnson, R.A., Murray, T.K., Ahmed, Z., et al. (2016). Imaging the accumulation and suppression of tau pathology using multiparametric MRI. Neurobiol. Aging 39, 184–194.

Holtmaat, A., Bonhoeffer, T., Chow, D.K., Chuckowree, J., De Paola, V., Hofer, S.B., Hübener, M., Keck, T., Knott, G., Lee, W.-C.A., et al. (2009). Long-term, high-resolution imaging in the mouse neocortex through a chronic cranial window. Nat. Protoc. 4, 1128–1144.

Hoover, B.R., Reed, M.N., Su, J., Penrod, R.D., Kotilinek, L.A., Grant, M.K., Pitstick, R., Carlson, G.A., Lanier, L.M., Yuan, L.-L., et al. (2010). Tau Mislocalization to Dendritic Spines Mediates Synaptic Dysfunction Independently of Neurodegeneration. Neuron 68, 1067–1081.

Hunsberger, H.C., Rudy, C.C., Weitzner, D.S., Zhang, C., Tosto, D.E., Knowlan, K., Xu, Y., and Reed, M.N. (2014). Effect size of memory deficits in mice with adult-onset P301L tau expression. Behav. Brain Res. 272, 181–195.

Hunsberger, H.C., Rudy, C.C., Batten, S.R., Gerhardt, G.A., and Reed, M.N. (2015). P301L tau expression affects glutamate release and clearance in the hippocampal trisynaptic pathway. J. Neurochem. 132, 169–182.

Ittner, A., and Ittner, L.M. (2018). Dendritic Tau in Alzheimer’s Disease. Neuron 99, 13–27.

Ittner, L.M., Ke, Y.D., Delerue, F., Bi, M., Gladbach, A., Eersel, J. van, Wölfing, H., Chieng, B.C., Christie, M.J., Napier, I.A., et al. (2010). Dendritic Function of Tau Mediates Amyloid-β Toxicity in Alzheimer’s Disease Mouse Models. Cell 142, 387–397.

Jackson, J., Jambrina, E., Li, J., Marston, H., Menzies, F., Phillips, K., and Gilmour, G. (2019). Targeting the Synapse in Alzheimer’s Disease. Front. Neurosci. 13.

Jackson, J.S., Witton, J., Johnson, J.D., Ahmed, Z., Ward, M., Randall, A.D., Hutton, M.L., Isaac, J.T., O’Neill, M.J., and Ashby, M.C. (2017). Altered Synapse Stability in the Early Stages of Tauopathy. Cell Rep. 18, 3063–3068.

Kneynsberg, A., Combs, B., Christensen, K., Morfini, G., and Kanaan, N.M. (2017). Axonal Degeneration in Tauopathies: Disease Relevance and Underlying Mechanisms. Front. Neurosci. 11.

Kopeikina, K.J., Polydoro, M., Tai, H.-C., Yaeger, E., Carlson, G.A., Pitstick, R., Hyman, B.T., and Spires-Jones, T.L. (2013). Synaptic alterations in the rTg4510 mouse model of tauopathy. J. Comp. Neurol. 521, 1334–1353.

Lee, J.-E., and Han, P.-L. (2013). An Update of Animal Models of Alzheimer Disease with a Reevaluation of Plaque Depositions. Exp. Neurobiol. 22, 84–95.

Menkes-Caspi, N., Yamin, H.G., Kellner, V., Spires-Jones, T.L., Cohen, D., and Stern, E.A. (2015). Pathological Tau Disrupts Ongoing Network Activity. Neuron 85, 959–966.

Miyamoto, T., Stein, L., Thomas, R., Djukic, B., Taneja, P., Knox, J., Vossel, K., and Mucke, L. (2017). Phosphorylation of tau at Y18, but not tau-fyn binding, is required for tau to modulate NMDA receptor-dependent excitotoxicity in primary neuronal culture. Mol. Neurodegener. 12, 41.

Mondragón-Rodríguez, S., Trillaud-Doppia, E., Dudilot, A., Bourgeois, C., Lauzon, M., Leclerc, N., and Boehm, J. (2012). Interaction of Endogenous Tau Protein with Synaptic Proteins Is Regulated by N-Methyl-d-aspartate Receptor-dependent Tau Phosphorylation. J. Biol. Chem. 287, 32040–32053.

Mudher, A., Colin, M., Dujardin, S., Medina, M., Dewachter, I., Alavi Naini, S.M., Mandelkow, E.-M., Mandelkow, E., Buée, L., Goedert, M., et al. (2017). What is the evidence that tau pathology spreads through prion-like propagation? Acta Neuropathol. Commun. 5, 99.

Nelson, P.T., Alafuzoff, I., Bigio, E.H., Bouras, C., Braak, H., Cairns, N.J., Castellani, R.J., Crain, B.J., Davies, P., Del Tredici, K., et al. (2012). Correlation of Alzheimer disease neuropathologic changes with cognitive status: a review of the literature. J. Neuropathol. Exp. Neurol. 71, 362–381.

Pickett, E.K., Henstridge, C.M., Allison, E., Pitstick, R., Pooler, A., Wegmann, S., Carlson, G., Hyman, B.T., and Spires-Jones, T.L. (2017). Spread of tau down neural circuits precedes synapse and neuronal loss in the rTgTauEC mouse model of early Alzheimer’s disease. Synapse 71, n/a-n/a.

Polydoro, M., Acker, C.M., Duff, K., Castillo, P.E., and Davies, P. (2009). Age-Dependent Impairment of Cognitive and Synaptic Function in the htau Mouse Model of Tau Pathology. J. Neurosci. 29, 10741–10749.

Polydoro, M., Dzhala, V.I., Pooler, A.M., Nicholls, S.B., McKinney, A.P., Sanchez, L., Pitstick, R., Carlson, G.A., Staley, K.J., Spires-Jones, T.L., et al. (2014). Soluble pathological tau in the entorhinal cortex leads to presynaptic deficits in an early Alzheimer’s disease model. Acta Neuropathol. (Berl.) 127, 257–270.

Ramsden, M., Kotilinek, L., Forster, C., Paulson, J., McGowan, E., SantaCruz, K., Guimaraes, A., Yue, M., Lewis, J., Carlson, G., et al. (2005). Age-dependent neurofibrillary tangle formation, neuron loss, and memory impairment in a mouse model of human tauopathy (P301L). J. Neurosci. Off. J. Soc. Neurosci. 25, 10637–10647.

van Rij, J., Hendriks, P., van Rijn, H., Baayen, R.H., and Wood, S.N. (2019). Analyzing the Time Course of Pupillometric Data. Trends Hear. 23, 2331216519832483.

Rocher, A.B., Crimins, J.L., Amatrudo, J.M., Kinson, M.S., Todd-Brown, M.A., Lewis, J., and Luebke, J.I. (2010). Structural and functional changes in tau mutant mice neurons are not linked to the presence of NFTs. Exp. Neurol. 223, 385–393.

Rosenmann, H., Grigoriadis, N., Eldar-Levy, H., Avital, A., Rozenstein, L., Touloumi, O., Behar, L., Ben-Hur, T., Avraham, Y., Berry, E., et al. (2008). A novel transgenic mouse expressing double mutant tau driven by its natural promoter exhibits tauopathy characteristics. Exp. Neurol. 212, 71–84.

Santacruz, K., Lewis, J., Spires, T., Paulson, J., Kotilinek, L., Ingelsson, M., Guimaraes, A., DeTure, M., Ramsden, M., McGowan, E., et al. (2005). Tau suppression in a neurodegenerative mouse model improves memory function. Science 309, 476–481.

Scheff, S.W., Price, D.A., Schmitt, F.A., and Mufson, E.J. (2006). Hippocampal synaptic loss in early Alzheimer’s disease and mild cognitive impairment. Neurobiol. Aging 27, 1372–1384.

Selkoe, D.J. (2002). Alzheimer’s disease is a synaptic failure. Science 298, 789–791.

Shadish, W.R., Zuur, A.F., and Sullivan, K.J. (2014). Using generalized additive (mixed) models to analyze single case designs. J. Sch. Psychol. 52, 149–178.

Spires-Jones, T.L., and Hyman, B.T. (2014). The Intersection of Amyloid Beta and Tau at Synapses in Alzheimer’s Disease. Neuron 82, 756–771.

Stephen, T.-L., Tamagnini, F., Piegsa, J., Sung, K., Harvey, J., Oliver-Evans, A., Murray, T.K., Ahmed, Z., Hutton, M.L., Randall, A., et al. (2019). Imbalance in the response of pre- and post-synaptic components to amyloidopathy. Sci. Rep. 9, 1–11.

Tackenberg, C., and Brandt, R. (2009). Divergent Pathways Mediate Spine Alterations and Cell Death Induced by Amyloid-β, Wild-Type Tau, and R406W Tau. J. Neurosci. 29, 14439–14450.

Tai, H.-C., Serrano-Pozo, A., Hashimoto, T., Frosch, M.P., Spires-Jones, T.L., and Hyman, B.T. (2012). The synaptic accumulation of hyperphosphorylated tau oligomers in Alzheimer disease is associated with dysfunction of the ubiquitin-proteasome system. Am. J. Pathol. 181, 1426–1435.

Thevenaz, P., Ruttimann, U.E., and Unser, M. (1998). A pyramid approach to subpixel registration based on intensity. IEEE Trans. Image Process. 7, 27–41.

Thies, E., and Mandelkow, E.-M. (2007). Missorting of Tau in Neurons Causes Degeneration of Synapses That Can Be Rescued by the Kinase MARK2/Par-1. J. Neurosci. 27, 2896–2907.

Wang, X., Smith, K., Pearson, M., Hughes, A., Cosden, M.L., Marcus, J., Hess, J.F., Savage, M.J., Rosahl, T., Smith, S.M., et al. (2018). Early intervention of tau pathology prevents behavioral changes in the rTg4510 mouse model of tauopathy. PLOS ONE 13, e0195486.

Wood, S.N. (2017). Generalized additive models: an introduction with R (CRC Press).

Yang, D. (2013). Perinatal Suppression of Tau P301L Has a Long Lasting Preventive Effect against Neurodegeneration. Int. J. Neuropathol. 53–59.

Yoshiyama, Y., Higuchi, M., Zhang, B., Huang, S.-M., Iwata, N., Saido, T.C., Maeda, J., Suhara, T., Trojanowski, J.Q., and Lee, V.M.-Y. (2007). Synapse loss and microglial activation precede tangles in a P301S tauopathy mouse model. Neuron 53, 337–351.

Zempel, H., and Mandelkow, E. (2014). Lost after translation: missorting of Tau protein and consequences for Alzheimer disease. Trends Neurosci. 37, 721–732.

Zhou, L., McInnes, J., Wierda, K., Holt, M., Herrmann, A.G., Jackson, R.J., Wang, Y.-C., Swerts, J., Beyens, J., Miskiewicz, K., et al. (2017). Tau association with synaptic vesicles causes presynaptic dysfunction. Nat. Commun. 8, 15295.

